# Olfactory coding in the turbulent realm

**DOI:** 10.1101/179085

**Authors:** Vincent Jacob, Christelle Monsempès, Jean-Pierre Rospars, Jean-Baptiste Masson, Philippe Lucas

**Affiliations:** Institute of Ecology and Environmental Sciences, INRA, route de St Cyr, 78000 Versailles, France; Peuplements végétaux et bioagresseurs en milieu végétal, CIRAD, Université de la Réunion, 97715 Saint Pierre, Ile de la Réunion, France; Decision and Bayesian Computation, Pasteur Institute, 25-28 rue du Dr Roux, 75015 Paris, France; Bioinformatics and Biostatistics Hub, C3BI, Pasteur Institute, CNRS, 25-28 rue du Dr Roux, 75015 Paris, France

## Abstract

Long-distance olfactory search behaviors depend on odor detection dynamics. Due to turbulence, olfactory signals travel as bursts of variable concentration and spacing and are characterized by long-tail distributions of odor/no-odor events, challenging the computing capacities of olfactory systems. How animals encode complex olfactory scenes to track the plume far from the source remains unclear. Here we focus on the coding of the plume temporal dynamics in moths. We compare responses of olfactory receptor neurons (ORNs) and antennal lobe projection neurons (PNs) to sequences of pheromone stimuli either with white-noise patterns or with realistic turbulent temporal structures simulating a large range of distances (8 to 64 m) from the odor source. For the first time, we analyze what information is extracted by the olfactory system at large distances from the source. Neuronal responses are analyzed using linear–nonlinear models fitted with white-noise stimuli and used for predicting responses to turbulent stimuli. We found that neuronal firing rate is less correlated with the dynamic odor time course when distance to the source increases because of improper coding during long odor and no-odor events that characterize large distances. Rapid adaptation during long puffs does not preclude however the detection of puff transitions in PNs. Individual PNs but not individual ORNs encode the onset and offset of odor puffs for any temporal structure of stimuli. A higher spontaneous firing rate coupled to an inhibition phase at the end of PN responses contributes to this coding property. This allows PNs to decode the temporal structure of the odor plume at any distance to the source, an essential piece of information moths can use in their tracking behavior.

**Author Summary:** Long-distance olfactory search is a difficult task because atmospheric turbulence erases global gradients and makes the plume discontinuous. The dynamics of odor detections is the sole information about the position of the source. Male moths successfully track female pheromone plumes at large distances. Here we show that the moth olfactory system encodes olfactory scenes simulating variable distances from the odor source by characterizing puff onsets and offsets. A single projection neuron is sufficient to provide an accurate representation of the dynamic pheromone time course at any distance to the source while this information seems to be encoded at the population level in olfactory receptor neurons.

## Introduction

A primary goal of olfaction research is to understand how complex olfactory scenes that occur in natural environment are processed by the olfactory system, in particular for orientation behaviors to odor sources. However, until recently natural odor stimuli were not described quantitatively. Hence, most studies of olfactory physiology are restricted to static and white-noise stimuli, limiting our understanding of dynamic olfactory coding. Although realistic olfactory input signals were used to analyze olfactory coding [1-3], they were uncontrolled, so preventing to explore specific features of the olfactory signals and large distances to the odor source. Only a recent paper generated naturalistic odor plumes and described how adaption from olfactory receptor neurons (ORNs) to stimulus mean and variance contribute to encode intermittent odor stimuli [4]. Here we pursue a novel approach to the study of olfactory coding.

For most animals larger than a millimeter, odor cue transport is dominated by turbulence [5-7]. Turbulence is thus a major key to understand animal orientation behavior from olfactory coding to decision dynamics. It prevents formation of gradients pointing towards the source and structures the plume in isolated odor patches that deform spatially as they are transported by the wind. Thus, odor detection is discontinuous. In turbulent conditions, temporal dynamics of odor/no-odor detection is dominated by long-tail statistics [8]. Furthermore, these statistics change with the distance to the odor source. The probability distributions for the duration and spacing in time of odor filaments become more heterogeneous with more short and long odor/no odor events when the distance increases [8]. Therefore, in real environments statistics of odor detection strongly differ from a white-noise pattern and are not characterized by a main frequency. For example, far from the source (∼100 m) a detection provides limited information on the time and duration of the next detection.

Male moths tracking plumes of sex pheromone released by their conspecific females is a paradigmatic example of olfactory searches [9, 10]. Successful searches are achieved with starting points at tens to hundreds of meters from the source [11-13]. The statistics of intermittent odor signals play a major role in the dynamics of moth tracking behaviors. Timing is also an important information for the vertebrate olfactory system [14-19]. However, opposite to vertebrates the insect nose is everted and can continuously monitor odor stimuli whose temporal structure is not distorted by respiratory cycles [18-21] and differential sorption into the mucus layer of the olfactory epithelium [22-25]. The rapid responses and rich dictionary of maneuvers of moths in turbulent flows suggest efficient coding of olfactory plume dynamics [26]. In turbulent environments, the flight is empirically described by two dynamical behavioral features: upwind surge, associated to pheromone perception, and zigzagging, a cast maneuver to retrieve odor plumes when lost [27-29]. These features are correlated to odor detection dynamics. Artificial homogenous pheromone clouds, thus unrealistic compared to real environment, lead to inefficient searches [30-33]. Change in the temporal characteristic of odor encounters has immediate effects on the searching dynamics [29, 32, 34-37] and can have more effect than a 1000-fold change in pheromone concentration [38].

Sensory systems efficiently process the stimuli encountered in their natural environment, according to the efficient coding hypothesis [39]. Adaptation of neural coding to the statistics of inputs [40-42] is essential for understanding evolutionary forces driving the properties of nervous systems. Natural stimulations are frequently reported to have broadband distributions with power-law statistics (e.g. for vision [43]). Olfactory signals involve the complex properties of turbulence and their dynamics are characterized by their extreme heterogeneity, long-tail statistics and thus multi-scale variability. In that sense, the olfactory dynamics encountered by individual moths can be “far” from an average statistics induced by the environment. This creates significant challenges in order to properly code the signal. An optimal olfactory coding should be able to reliably encode the plume temporal structure at all relevant distances from the source. Essential characteristics of the puffs-blanks dynamics of the signal required to reach an olfactory source remain unknown. Analyses from experimental trajectories [9, 44] are often performed without knowing the precise olfactory signal (as it is inaccessible). As a consequence, theoretical analyses of orientation strategies have to be performed without knowing the computing capacities of insects.

In this work, we use the quantitative analysis of plume dynamics in turbulent conditions [8] to study moths’ neuronal coding of olfactory plumes with realistic turbulent dynamics and investigate for the first time what information is extracted at large distances from the source. We focus on how ORNs and antennal lobe projection neurons (PNs) encode the temporal characteristics of the pheromone plume in the Noctuid moth *Agrotis ipsilon*. We show that firing activities of ORNs and PNs are less correlated with the temporal structure of olfactory scenes when they simulate increasing distances to the source. Individual PNs but not individual ORNs encode puff onsets and puff offsets at multiple temporal scales. Thus, PNs can reconstruct the odor sequence at any distance from the source.

## Results

### Linear-nonlinear (L-N) model fits neuronal response to white-noise stimuli

In order to study the neuronal coding of dynamic pheromone stimuli in the moth olfactory pathway, we recorded the firing rate of individual pheromone-sensitive ORNs and PNs in whole-insect preparations while presenting time-varying binary sequences of pheromone stimuli. We show the spiking activity of an ORN (Fig 1A-C) and a PN (Fig 1D-F) in response to a binary sequence of white-noise stimuli. We observed a good match between stimulus puffs and spike trains. A total of 209 ORNs and 97 PNs were recorded while presenting long duration (15 min and 5 min, respectively) white-noise sequences of pheromone stimuli.

**Figure 1.**
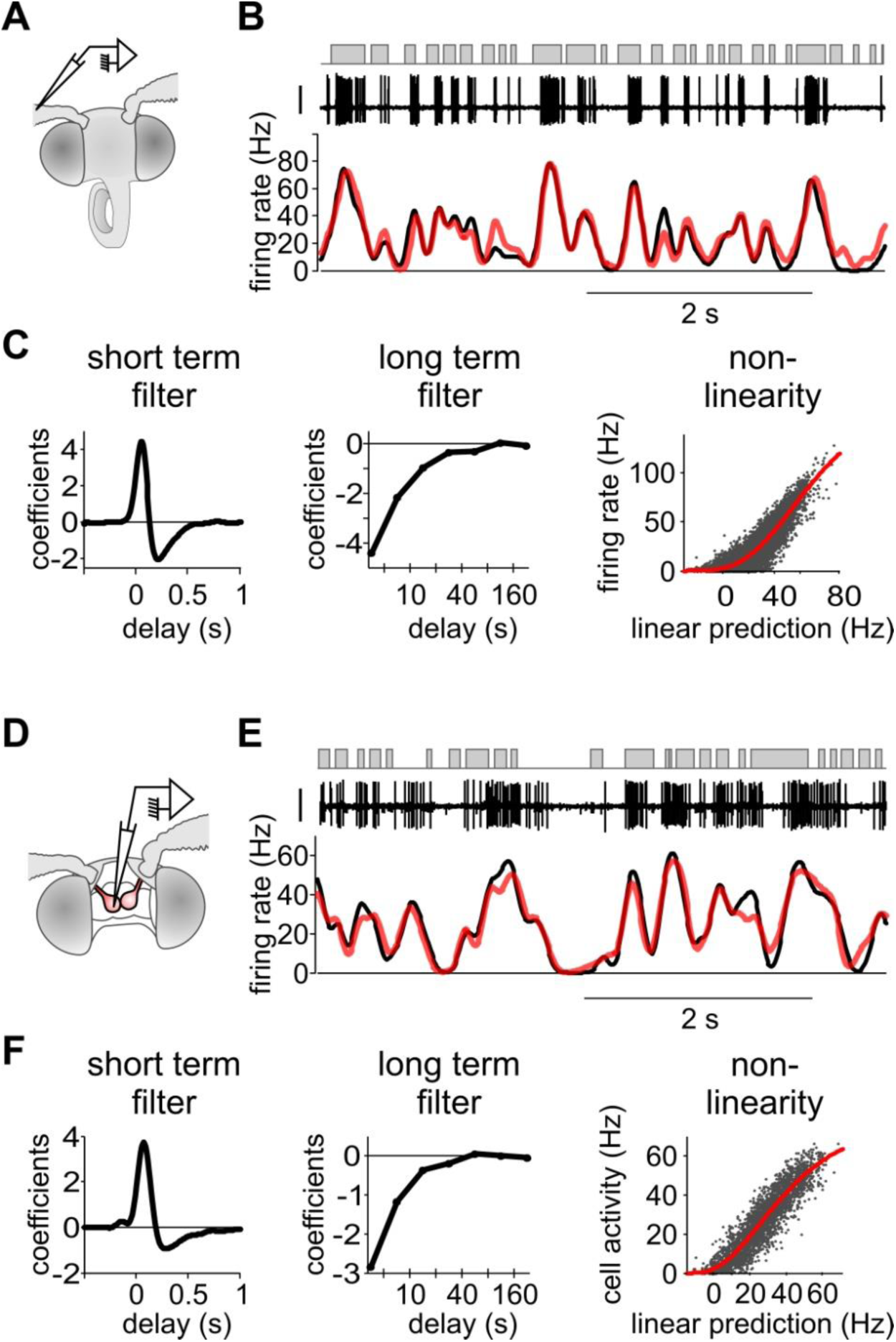
L-N model captures the responses of both ORNs and ALNs to white-noise stimuli. (A) Moth preparation for ORN recording. A tungsten electrode is inserted at the basis of a sensillum. (B) Response of an ORN to white-noise odor stimuli (correlation time 20 ms). Top: stimulus sequence (puffs in grey). Middle: action potential recordings (vertical bar 1 mV). Bottom: firing rate dynamic calculated with a Gaussian kernel (black) and predicted with an L-N model (red). (C) L-N model fitted to the response of the same neuron as in (B) to 15 min of white-noise stimuli. Left to right: short term linear filter, long term linear filter calculated on a log time scale, static nonlinearity (grey dots: measured vs predicted values, red line: fitted Hill function). (D) Moth preparation for PN recording. A glass electrode is inserted in the cumulus of an antennal lobe. (D-F) Same as (B-C) but on a PN.

In order to characterize the temporal properties of neural coding of olfactory stimuli we fitted a L-N model to the firing activity of each neuron (Fig 1C,F) consisting of a linear filter combined with a nonlinear function. First, we calculated the linear filters (or transfer functions) between white-noise sequences and the instantaneous firing rate with Lasso regularization on short- and long-term components. The linear filters were subdivided in two different time scales: a short-term component (time window −0.5 to 2.5 s) prolonged by a long-term component (2.5 to 320s) with exponential time sampling size. The long-term component takes into account the decrease of evoked firing activity during the stimulation sequence (time constant of the exponential decay is 89 s for ORNs and 15 s for PNs, Fig 2A). The neuronal responsiveness was restored after 3 minutes without stimuli both in ORNs (Fig 2B) and PNs (Fig 2C). The slow decrease of activity is thus a physiological process and did not result from a degradation of the quality of recordings or from a depletion of pheromone in the stimulus pipette. Long-term filters weighted 10.4% for ORNs (median value; bootstrap confidence interval: 9.9 to 12%) and 8.9% for PNs (median value; bootstrap confidence interval: 7 to 10.1%) of the total linear filter. Predictions of firing rates using linear filters were frequently negative indicating discrepancies between the measured neuronal activities and their linear prediction. This was expected since neurons are known to be nonlinear integrators of their inputs. Second, a simple static nonlinearity was used to correct the linear prediction above. We estimated the static nonlinearity with fitting a Hill function between linear prediction and measured activity (sigmoid curve, Fig 1C,F). Measured and L-N model-predicted neuronal activities are superimposed for the two example neurons (Fig 1B,E). Model predictions reproduced most of the fluctuations of the measured firing rates.

**Figure 2.**
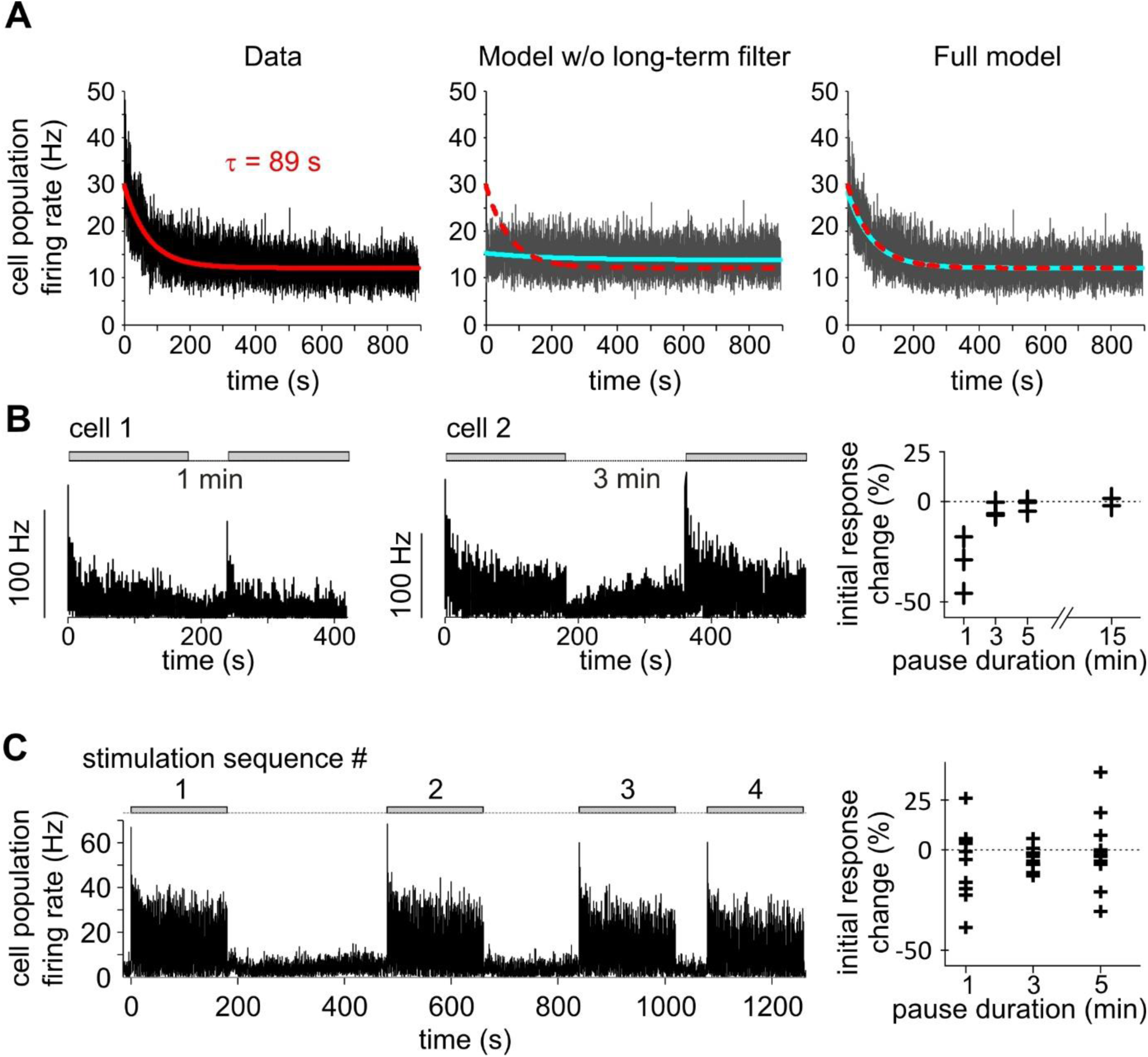
A reversible long term adaptation to dynamical stimulations is only captured with a long-term filter. (A) ORN responses to long duration white-noise stimuli and L-N model fits. Firing rate measured in cell population (left), predicted by L-N models without long-term filter (middle) and predicted by the full L-N model used in the paper (right). Red and dotted red: exponential curve fitted to the data, blue: exponential curves fitted to the model prediction. (B) The slow adaptation is reversible. Left: example of ORN response to two 3-min-long noise sequences (grey area) interleaved by a 1-min-long pause. The initial response (maximal firing rate in the first 5 s) to the beginning of the second noise sequence is smaller. Middle: example of ORN response to two 3-min-long noise sequences interleaved by a 3-min-long pause. The initial response to the second noise sequence is similar to the initial response to the first sequence. Right: initial response changes in ORNs depend on the pause duration (cell population = 11). (C) PN population response to 4 consecutive frozen noise sequences (n = 16 cells). The noise sequence (grey areas) was 3-min long and its repetitions were interleaved by respectively 5, 3 and 1 min pauses. Right: initial response changes in PNs (n = 10).

We quantified the performance of the L-N model with the coefficient of determination *R*^2^ calculated between the L-N prediction and the measured firing rate. We tested different types of white-noise sequences varying in the pheromone dose (only in ORNs) and in the correlation time (in ORNs and PNs). In ORNs, *R*^2^ increased significantly with the pheromone dose (ANOVA, p<0.001), revealing that the relationship between firing rate and stimuli is tighter at high pheromone concentrations (Fig 3A). Furthermore, *R*^2^ was lower for the fastest white-noise sequences (time step = 10 ms) both in ORNs (ANOVA, p<0.001; Fig 3A) and in PNs (Wilcoxon’s rank sum test for independent recordings, p<0.01; Wilcoxon’s signed rank test for paired samples p<0.05, n = 15; Fig 3B-C). Finally, we observed that some PNs did not encode well the temporal fluctuations of the white-noise. Some of these PNs might have been damaged by the dissection and/or when introducing the recording electrode in the antennal lobe. However, we previously described in *A. ipsilon* the heterogeneity in sensitivity and delay of response across PNs from the cumulus [45]. Therefore, some PNs with low quality L-N prediction may correspond to weakly sensitive neurons or to neurons that do not encode well the temporal dynamics of stimuli and better encode other properties of the stimulus such as its quality or quantity. In the following analysis of responses to turbulent stimuli we considered only the PNs with *R*^2^ > 0.3 (70 out of 97 cells) for white-noise sequences.

**Figure 3.**
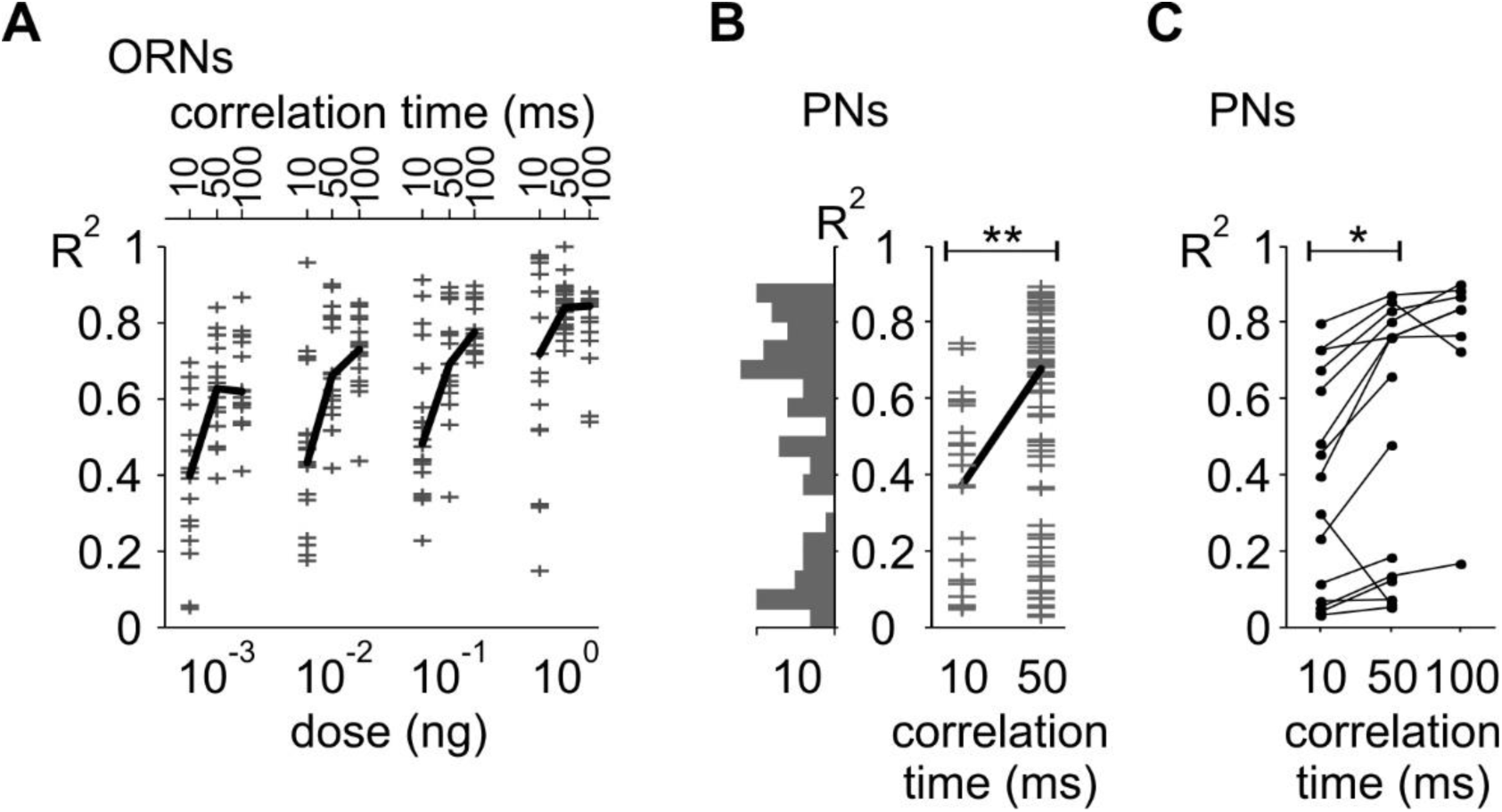
Performances of L-N model predictions. (A) Distribution of coefficient of determination *R*^2^ between model prediction and ORN responses to white-noise stimuli. Grey crosses: individual cells. Black segment: median values. *R*^2^ depends both on noise correlation time (ANOVA, p<10^-10^) and pheromone load (ANOVA, p<10^-10^). (B) Distribution of *R*^2^ between model prediction and PN responses to noise stimuli, independent samples. Left: the distribution histogram shows the heterogeneity of *R*^2^ values. **: p<0.01 Wilcoxon’s rank sum test. (C) *R*^2^ between model prediction and PN responses to noise stimuli, paired samples. For this data set, two or three noise sequences with different correlation times were tested for each cell. Line segments join the *R*^2^ calculated for the same cell. *: p<0.05 Wilcoxon’s signed rank test.

### Olfactory neurons display a family of linear filter shapes that can be estimated with simple sets of parameters

The shape of a linear filter characterizes how a neuron integrates time-fluctuating stimuli. The shape of individual linear filters is remarkably homogeneous for ORNs and more diverse for PNs (Fig 4A-B). We analyzed the variability in shape of short-term filters with a principal component analysis as performed by Geffen et al. [46]. For ORNs, the first principal component (PC1) explains 79% of the filter variance (Fig 4C). It has a positive phase followed by a negative one and its normalized integrated area is close to 0 (-0.07). Hence, it corresponds to purely phasic responses to olfactory stimuli. The first two principal components (PC1 and PC2) together explain 93.5% of the filter variance; PC2 has essentially one positive phase and a normalized integrated area close to 1 (0.90). Thus, PC2 corresponds to tonic responses to olfactory stimuli. As PC1 and PC2 account for a disproportionate amount of the filter variance, each ORN linear filter can be considered as a linear combination of these two components. Individual linear filters were normalized in amplitude so that each filter corresponds to one dot lying on the *n*-sphere of radius 1 in the space of the *n* principal components. A projection of this representation on the 2-D space of PC1 and PC2 is shown (Fig 4E). Most of the dots lie close to the unit circle, indicating that the first two principal components are sufficient to describe the filter shape with the exception of a few ORNs stimulated at the highest pheromone concentration. The relative contribution of the first two principal components to the linear filter can be directly read as the angle *φ* on that circle. A filter with an angle *φ* = 0 means that responses were purely phasic, *φ* > 0 means that responses were phasic-tonic, and *φ* < 0 observed at high pheromone concentrations means that responses were phasic and then inhibitory. Few cases were close to tonic excitatory responses (*φ* = π/2).

**Figure 4.**
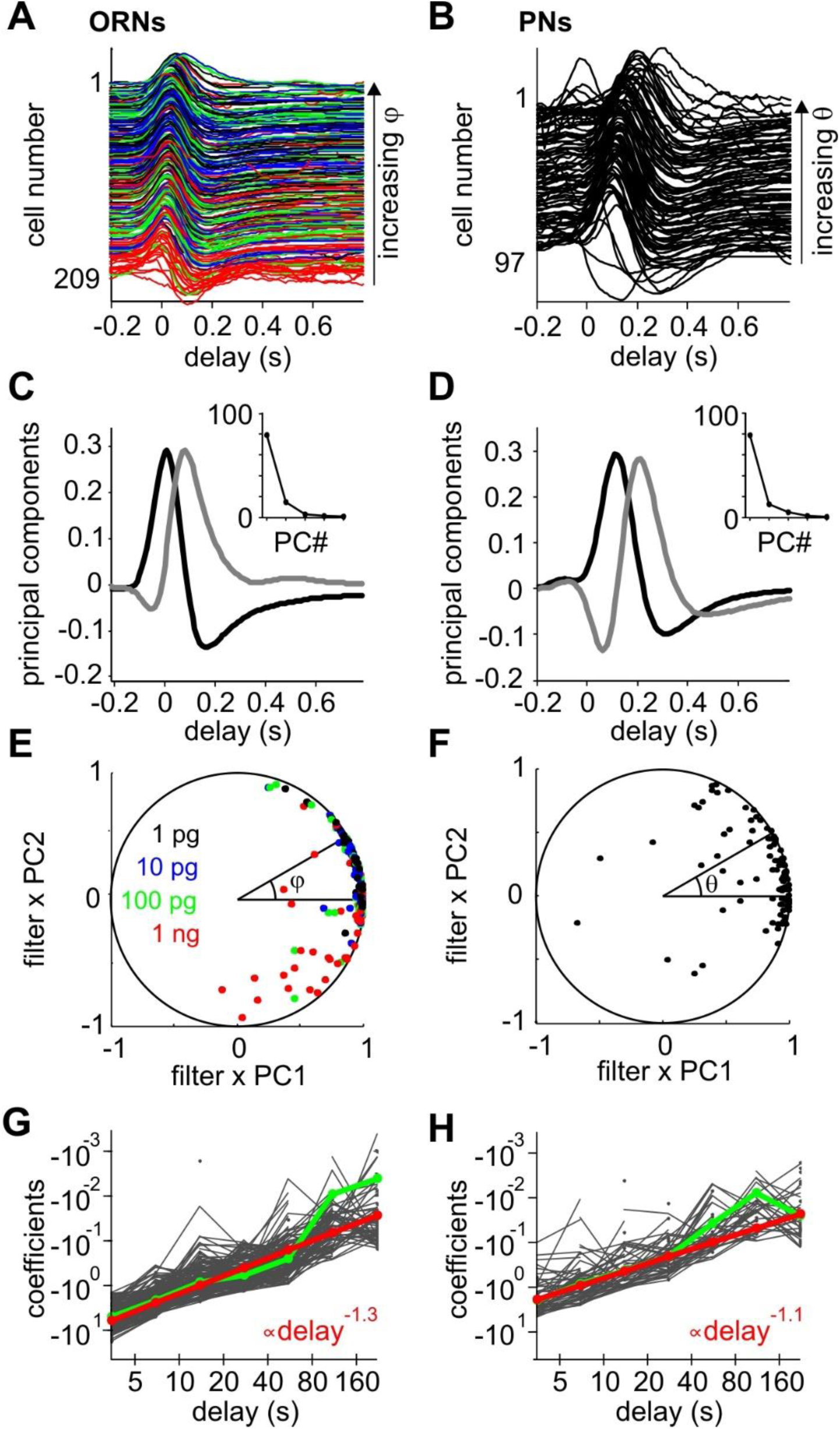
Parametrization of the shapes of linear filter sets. The diversity of short-term filters is analyzed as in Geffen et al. (2009). Analysis of the diversity of long-term filters is adapted from Pozzorini et al. (2013). (A) Individual short-term linear filters for 209 ORNs sorted by decreasing parameter *φ* (see (E)). Same color code for the different pheromone loads in the stimulation pipette as in (E). (B) Individual short term linear filters for 97 PNs, sorted by decreasing *θ* (see (F)). (C) First (black) and second (grey) principal components of the ORN short-term filter set. Inset: percentage of variance explained by the first 5 components. (D) First two principal components for the PN short-term filter set. Same conventions as in (C). The time shift of principal components for PNs when compared to ORNs reflects the difference of delay for the pheromone to reach the antennae with the odor delivery devices used for ORN and PN recordings. (E) Projection of normalized ORN short term filters on the subspace spanned by the first two components. If a filter is solely described by the two components, then the corresponding dot lies on the unit circle. Each short term filter can be parametrized by its angle *φ*. (F) Projection of normalized PN short term filters on the subspace spanned by the first two components. Each short term filter can be parametrized by its angle *θ*. (G) Individual ORN long term filters plotted on a log-log scale. Occasional values above 0, mostly for long delays, were not plotted. Green: median values. Red: power-law fit. (H) Individual PN long-term filters. Same conventions as in (G).

For PNs the same principal component analysis was performed (Fig 4D,F). PC1 explains 79% of the variance and PC1+PC2 92%. PC1 has a positive phase followed by a negative one resulting in a slightly positive normalized area (0.23). PC2 has a negative phase followed by a positive phase with a normalized area of 0.07. The contribution of the negative phase of PC2 cancels the positive phase of PC1 and therefore delays the response from stimulus onset. As a consequence the relative contribution of the first two components (angle *θ* on the unit circle) can be considered as the temporal phase lag between the neuronal response and the stimulation. Indeed as seen in Fig 4B, the position of the maximum of PN filters is delayed in time with increasing *θ*.

The individual long-term filters were distributed along a straight line on a log-log scale plot (Fig 4 G-H) both for ORNs and for PNs. We thus fitted a power-law function that revealed that both ORNs and PNs had similar scaling exponents (-1.3 for ORNs and −1.1 for PNs).

Linear filters were calculated for individual neurons in response to a sequence of white-noise stimuli. Only if it quantifies a stable cell property over time can it be used to predict neuronal response. We repeated up to 4 frozen sequences of white-noise in PNs interleaved by pauses of 1 to 5 minutes duration. The model was fitted to each sequence independently and the model coefficients were remarkably similar for the different responses of the same cell confirming the stability of linear filters (Fig 5). For PNs, the cell-specific linear filter calculated from white-noise stimuli was used to predict responses to turbulent stimuli in the same cell. However this was not possible for ORNs since responses to white-noise and turbulent stimuli were recorded from different cells. Instead we estimated individual linear filters with fitting three parameters: the angle *φ* of the short-term filter, the scaling exponent of the long-term filter *β* and the relative weight of long- and short-term filters *α*. The amplitude of the linear filter was not considered since the nonlinear function compensates for it.

**Figure 5.**
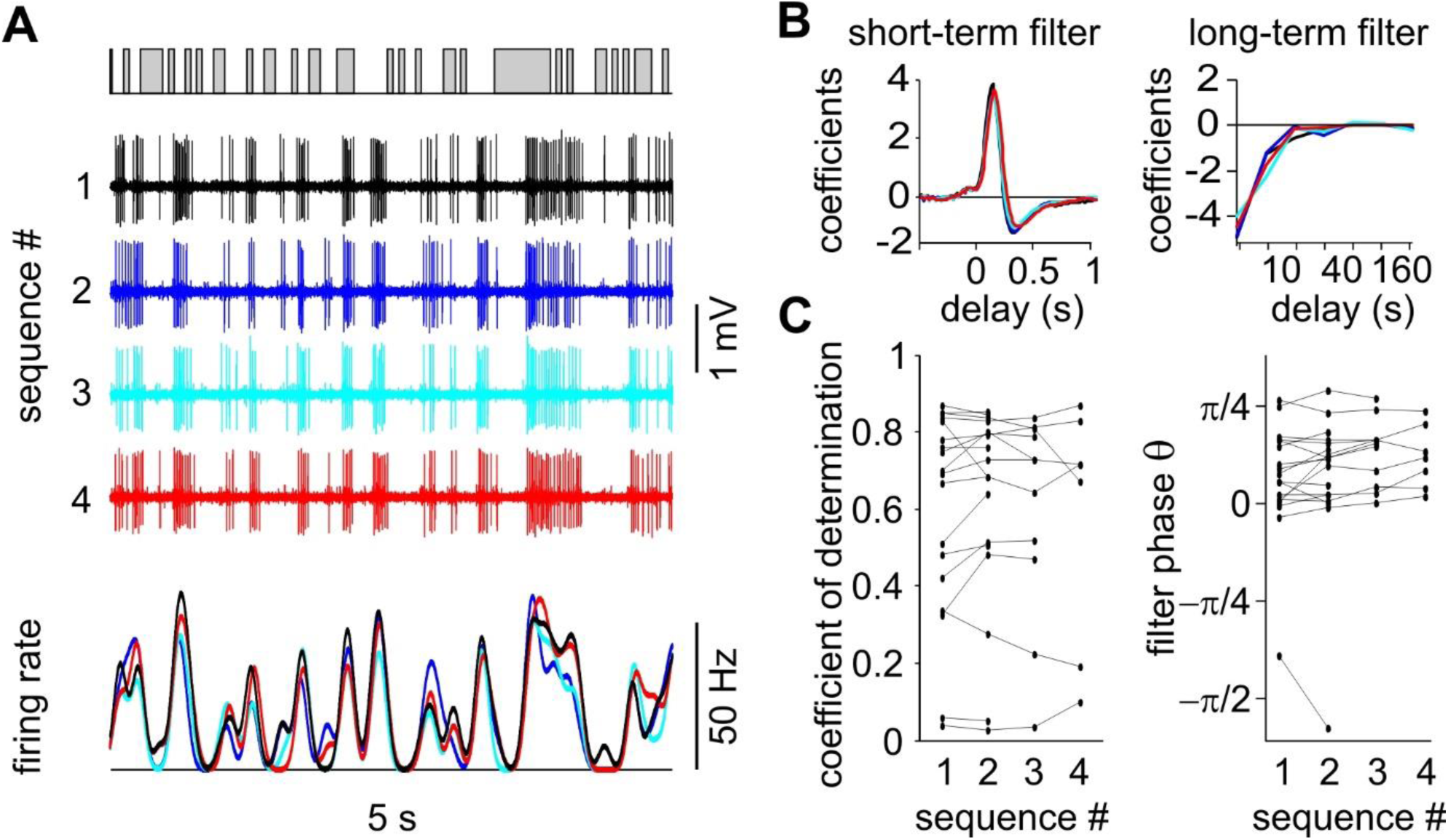
Stability of L-N model prediction for PNs. (A) Zoom on 4 consecutive responses from a PN to a frozen white-noise. Top: stimulation sequence. Middle: action potential recordings. Bottom: recorded firing rates. (B) Superimposed linear filters calculated independently for the same PN as in (A). (C) Distribution of the coefficient of determination and short-term filter phase for the 4 consecutive sequences. Same neuron values are joined by line segments. There was no significant change for any parameter (cell number = 18).

### Olfactory coding performance depends on the temporal structure of the pheromone plume

We stimulated the antenna with temporal sequences of pheromone stimuli having a temporal structure characteristic of turbulent environments. Turbulent sequences were generated as previously described [8] and consisted in binary sequences of pheromone stimuli with puffs of pheromone and clean air (blanks) between puffs of various durations (Fig 6A). The distribution of the duration of puffs and blanks in these sequences varied with the simulated distance from the odor source (Fig 6B). The distributions were more homogeneous at short distances from the source (at 8 m, 0.1-99.9% quantiles ranged from 126 ms to 8 s for puffs and from 126 ms to 12 s for blanks) and were broader at long distances with both shorter and longer events (at 64 m, 0.1-99.9% quantiles ranged from 16 ms to 22 s for puffs and from 16 ms to 89 s for blanks). These were characteristic long-tail distributions. For comparison we also show the distribution of the white-noise sequences used to fit the L-N model (Fig 6C). Durations of puffs and blanks have an exponential distribution in white-noise stimuli that is more homogeneous (for a correlation time of 50 ms, 0.1-99.9% quantiles ranged from 50 to 550 ms both for puffs and blanks) than the long-tail distributions in turbulent sequences.

**Figure 6.**
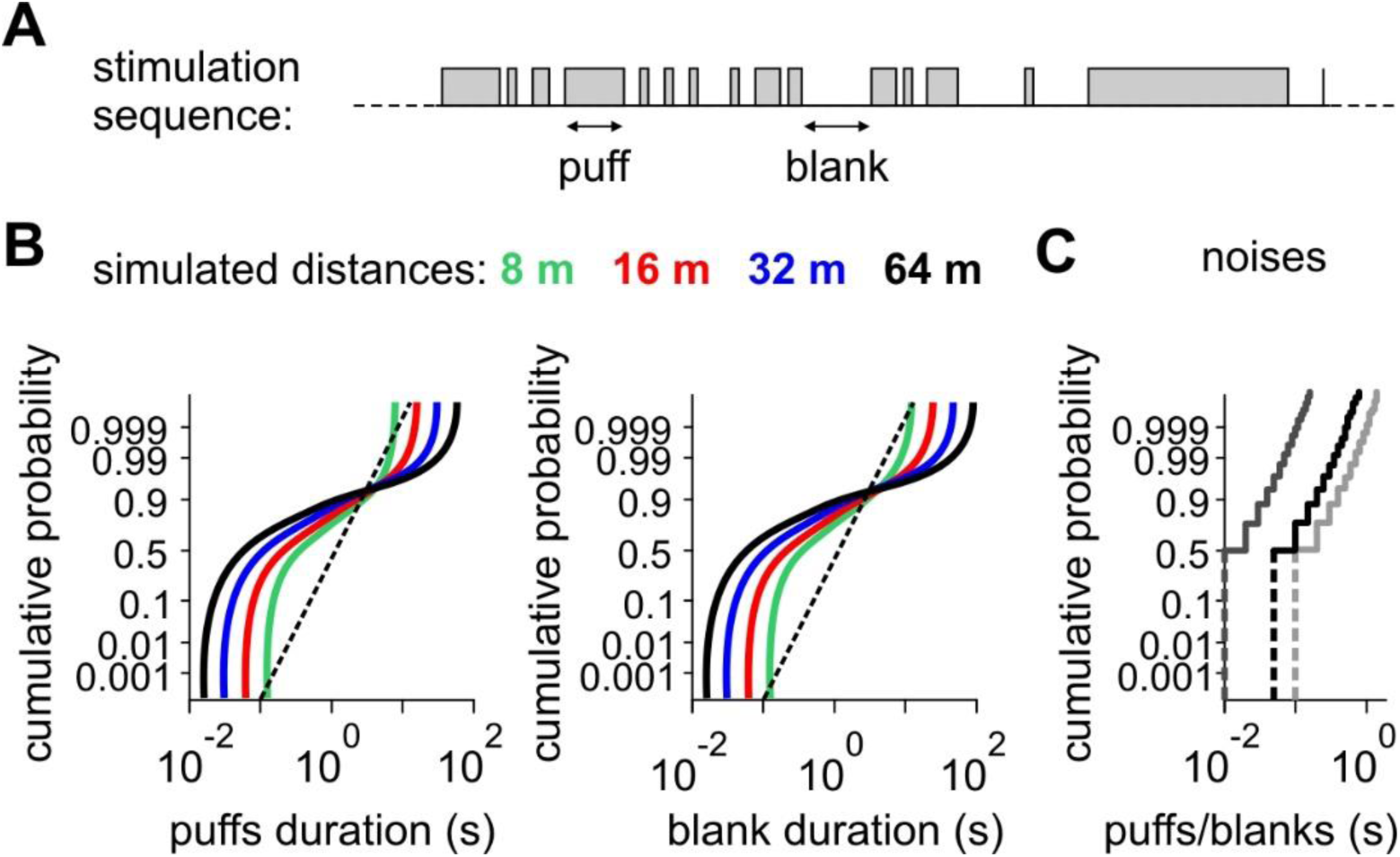
Turbulent stimuli have more events with extreme duration at long simulated distances.

(A) Example of a binary turbulent stimulation sequence. Puffs and blanks were defined as the presence and absence of pheromone displayed to the antenna. (B) Normal Probability Plot of the duration of puffs and blanks of turbulent stimuli. Colors code for the distance to the source. In Normal Probability Plots, any Gaussian distribution would appear as a straight line as shown (dotted line). In comparison turbulent stimuli have characteristic shape of long tail distributions with an excess of large and small values, and the excess increases with the distance from the source. (C) Normal Probability Plot for binary white-noises with different correlation time (dark grey, 10 ms; black, 50 ms; light grey, 100 ms). Duration of puffs and blanks have the same exponential distribution in white-noise, which by definition is not long-tail either.

L-N models obtained from binary white-noise stimuli were used to predict the response of ORNs and PNs to the turbulent stimuli. To correct statistical errors induced by nonlinearities [47], the non-linear part of the filter was re-estimated for the turbulent stimuli. Examples of responses and L-N predictions are shown in Fig 7A,D for two ORNs and one PN. We calculated the coefficients of determination *R*^2^ between model predictions and measured responses. In PNs the model fitted with the white-noise sequence predicts the response to turbulent stimuli (median *R*^2^ = 0.57; bootstrap confidence interval: 0.52 to 0.62). Removing any component of the model significantly decreased its performance in predicting responses (Table 1). These observations confirm the accuracy of the model and the critical role of the PC1/PC2 ratio.

**Table 1:**
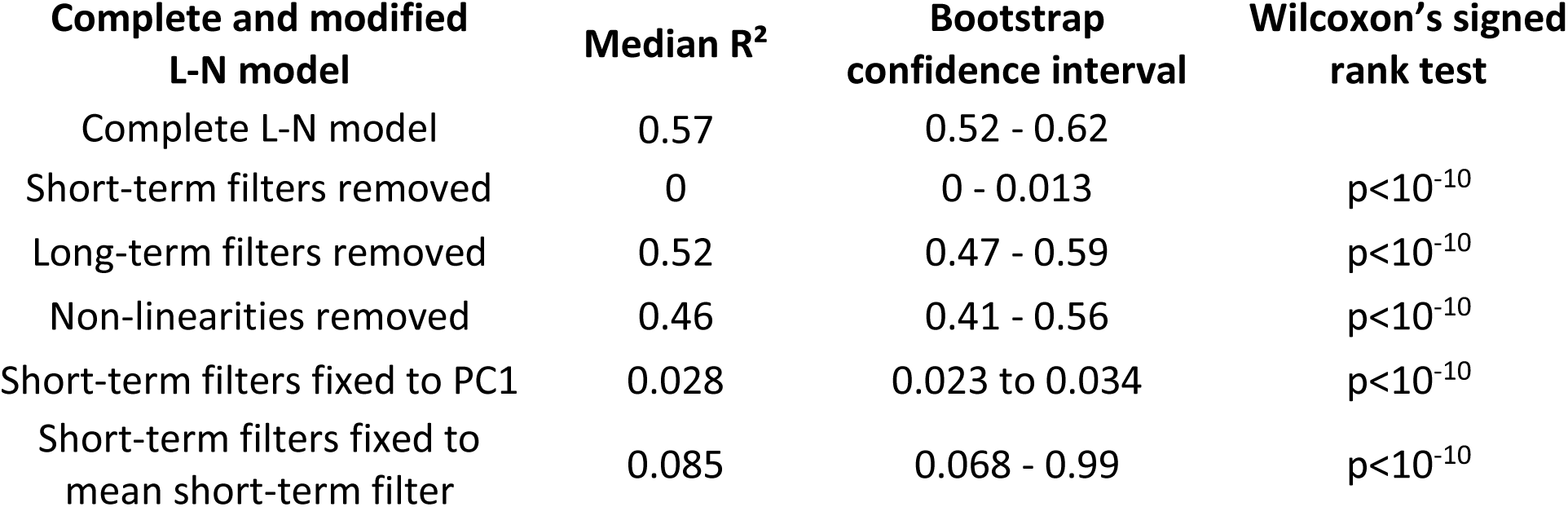
Coefficients of determination *R*^2^ between predictions and measured firing rates for the L-N model used in this work and several modified L-N models. All modification of the model significantly decreased its performance in predicting firing rates (Wilcoxon’s signed rank tests).

| Complete and modified L-N model | Median $R^2$ | Bootstrap confidence interval | Wilcoxon's signed rank test |
| --- | --- | --- | --- |
| Complete L-N model | 0.57 | 0.52 - 0.62 |  |
| Short-term filters removed | 0 | 0 - 0.013 | $p < 10^{-10}$ |
| Long-term filters removed | 0.52 | 0.47 - 0.59 | $p < 10^{-10}$ |
| Non-linearities removed | 0.46 | 0.41 - 0.56 | $p < 10^{-10}$ |
| Short-term filters fixed to PC1 | 0.028 | 0.023 to 0.034 | $p < 10^{-10}$ |
| Short-term filters fixed to mean short-term filter | 0.085 | 0.068 - 0.99 | $p < 10^{-10}$ |

**Figure 7.**
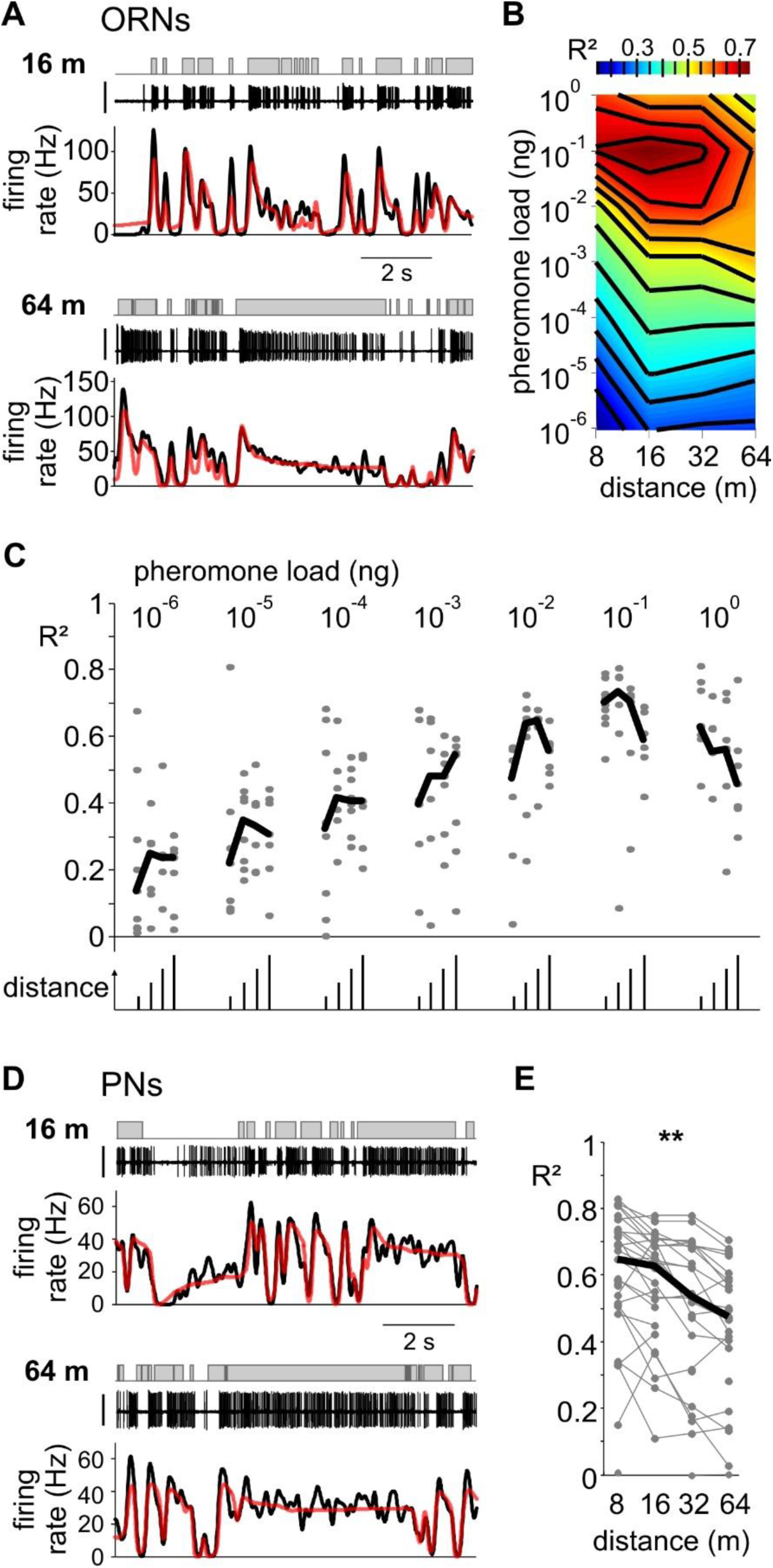
L-N model performance varies with the simulated distance during turbulent stimuli. (A) Response of two ORNs to turbulent stimuli simulating distances of 16 m (top) and 64 m (bottom). Same conventions as in Fig 1B. (B) Heatmap of the median coefficient of determination (*R*^2^) between L-N model prediction and measured ORN responses, plotted as a function of pheromone load and distance. Note that at high pheromone load, *R*^2^ is higher for short distances, but this is not true anymore for low pheromone loads. (C) Cell distribution of *R*^2^ between L-N prediction and measured ORN responses. Grey dots are *R*^2^ for individual cells, black segments show the median values. For each pheromone load value, the distance of the odorant source is 8, 16, 32 and 64 m from left to right. (D) Response of a PN to turbulent stimuli simulating distances of 16 m (top) and 64 m (bottom). Same conventions as in Fig 1B. (E) *R*^2^ between L-N prediction and measured PN response decreases with distance. One cell is represented by up to 4 dots joined by a grey line. Black line, median value. *R*^2^ decreased linearly with the log of the distance (**: p<0.01, permutation test applied on linear regression ramp).

Due to turbulence, variations of distances from the source during a moth olfactory search not only lead to complex changes in the whiff time dynamics, it also results in large and intricate fluctuations of pheromone concentrations within whiffs, i.e. peak concentration within whiffs does not decrease regularly with the distance to the source, so the highest concentrations are not necessarily all close to the source [8, 48]. Hence, to understand olfactory coding, we varied both the concentration and statistics of temporal dynamics for simulating different distances.

For ORNs, we tested turbulent stimuli simulating 4 different distances at pheromone concentrations over a large range encompassing 7 orders of magnitudes. The performance of the model clearly depended on pheromone concentration (ANOVA, p < 10^-10^, n = 192 cells) but did not depend on the simulated distance (ANOVA, p>0.3). More specifically, *R*^2^ increased linearly from 10^-6^ ng to 10^-1^ ng but for the highest concentration (1 ng) decreased back to the level of 10^-2^ ng (Fig 7C). This effect suggests that the range of concentrations we explored encompasses the physiological range. We represented this space with a phase diagram showing the dependence of olfactory coding performance (*R*^2^) on pheromone concentration (dose tested) and simulated distance (Fig 7B).

For each PN we tested up to 4 sequences of stimuli simulating different distances and presented in random order. To benefit from the paired structure of the data, we used specific statistics for testing the dependence of model performance on the temporal sequence. The coefficient of determination *R*^2^ was negatively correlated with the logarithm of the simulated distance (permutation test applied on linear regression ramp, p<10^-5^, n = 39 cells; Fig 7E). Olfactory coding performance in PNs depended on the temporal structure of the turbulent pheromone plume.

### Source of the divergence between modelled and measured firing rate during turbulent stimuli

We hypothesized that the lower ability of neurons to encode the olfactory signal when the distance is increased may reflect a difference in coding of the long puffs and blanks that are characteristic of large distances. To test this hypothesis we subdivided the responses to turbulent stimuli into different sub-regions of time that we categorized as: (a) onset of puffs (first 0.5 s), (b) tails of puffs (after 2 s from the beginning of puffs), (c) offset of puffs (first 0.5 s following the end of puffs lasting at least 0.1 s) and (d) tails of blanks (after 2 s from the end of puffs) (Fig 8A). As expected, ORN firing rate was the highest at the onset of puffs (median firing rate 19.8 Hz) and reached a steady state after prolonged exposure due to adaptation (7.3 Hz) (Fig 8B). A very low firing rate was observed in the absence of pheromone, both at the offset of puffs (0.02 Hz) and during tails of blanks (0.5 Hz). PN firing rate was the highest at the onset of puffs (median firing rate 21.2 Hz) and decreased during long puffs down to 18.5 Hz (Fig 8C). On average a transient inhibitory phase followed the end of puffs (1.9 Hz) and the firing rate rose up back to a higher level after a prolonged absence of pheromone (7.8 Hz).

**Figure 8.**
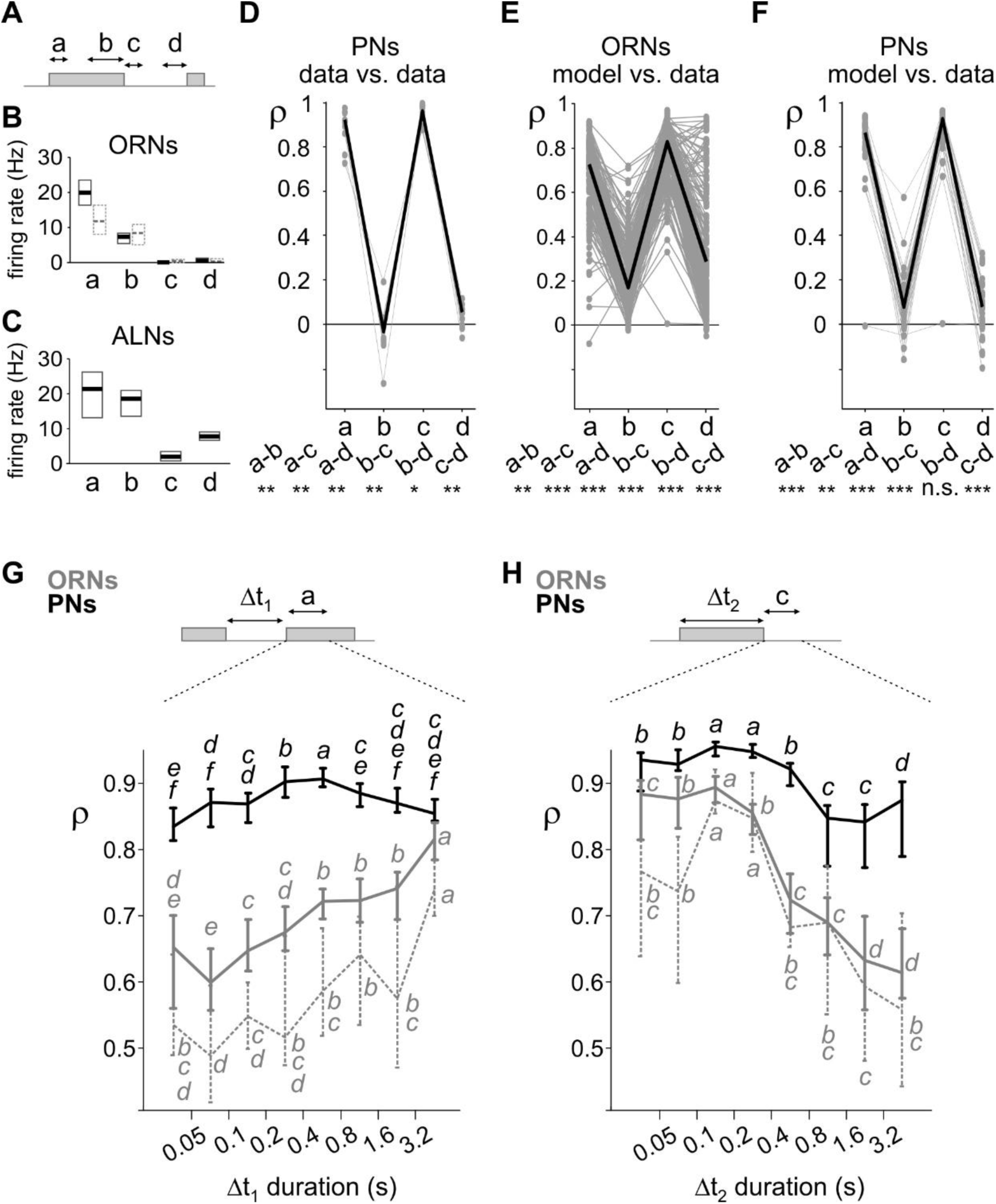
Neuronal coding varies with time sub-regions during turbulent stimuli. (A) Four different sub-regions of stimulus sequences were defined with (a) onset of puffs, (b) end of long puffs, (c) offset of puffs and (d) end of long blanks; see text for detailed definition. (B) (ORNs and in dotted grey the subpopulation of ORNs stimulated with pheromone loads of 10^-5^ and 10^-4^ ng) and (C) (PNs): median values and bootstrap confidence intervals (95%) of firing rate in the 4 sub-regions. (D) Correlation coefficient between consecutive responses of PNs to a sequence turbulent stimuli for the 4 sub-regions defined in (A). Distance was 16 m. Cell number = 13. For each cell, 4 grey dots joined by a grey line represent the median values of correlation coefficients *θ* during sub-regions. Black line joins the median values. Bottom: dual tests between sub-regions (*: p<0.05, **: p<0.01, ***: p<0.001, n.s.: non-significant, Wilcoxon’s signed rank tests). (E) Correlation coefficient between L-N prediction and ORN responses for the 4 sub-regions. Responses to all distances and doses are pulled together. Same conventions as in (D). (F) Correlation coefficient between L-N prediction and PN responses for the 4 sub-regions. Same conventions as in (D). (G) Correlation coefficient between L-N prediction and measured responses during the onset of puffs depends on the time delay from the end of the preceding puff. Plot shows median values and bootstrap confidence intervals. Grey: ORNs, dotted grey: subpopulation of ORNs stimulated with pheromone loads of 10^-5^ and 10^-4^ ng, black: PNs. Both for ORNs and for PNs, values with the same letters are not statistically different within a set of neurons (Wilcoxon’s signed rank test, p>0.05). (H) Correlation coefficient at the end of puffs depends on the duration of the preceding puff. Same convention as in (G).

To test how neurons encode the different time sub-regions, we repeated twice a frozen sequence of stimuli simulating a distance of 16 m on PNs (13 cells, 31 pairs of sequences, Fig 8D). We evaluated the correlation coefficient *ρ* between the measured responses to the first and second presentation of the stimulation sequence. *ρ* was calculated for all the time subregions then averaged for each category. It was larger at the onset (median *ρ* = 0.91) and offset of puffs (median *ρ* = 0.95), and indistinctively low both during tails of puffs (median *ρ* = and tails of blanks (median *ρ* = 0.05).

We then calculated the average correlation coefficient *ρ* between the L-N prediction and the measured firing rate. We observed the same patterns as for repeated measures: for ORNs *ρ* was maximal at the puff onset (median *ρ* = 0.72) and low during tails of puffs (median *ρ* = 0.17). At the offset of puffs *ρ* was high again (*ρ* = 0.82) and was low during tails of blanks (*ρ* = 0.28). All paired differences were significant with p<0.01 (paired Wilcoxon’s signed rank test, n = 192 cells; Fig 8E). The pattern of prediction performance was similar for PNs: the prediction quality was high at the onset (median *ρ* = 0.85) and at the offset of puffs (median *ρ* = 0.74) and low during tails of puffs (median *ρ* = 0.08) and tails of blanks (median *ρ* = 0.08). It is noteworthy that there was no difference in the quality of prediction of tails of puffs and tails of blanks (p>0.8, paired Wilcoxon’s signed rank test, n = 46 cells; Fig 8F).

This observation further confirms that PNs do not encode olfactory stimuli during prolonged presentation, but they encode well the beginning and the end of puffs which is sufficient to reconstruct the odor sequence.

### PNs but not ORNs encode olfactory stimuli at multiple time scales

We then focused on the two sub-regions with high correlation coefficients *ρ* (onset and offset of puffs) to know if the prediction performance varied with the duration of the preceding time interval. We observed a clear difference between ORNs and PNs. In ORNs the L-N model predicted well the beginning of isolated puffs, remote from any previous stimulation (time interval > 3.2 s). The prediction became gradually lower as the preceding puff was at a shorter time interval (Fig 8G). In addition, the offset of short puffs (duration < s) was better predicted than the offset of longer puffs (Fig 8H). Thus, the coding in ORNs appears adapted for a finite set of temporal profiles: short and isolated puffs. These statistical characteristics are associated to close distance dynamics to the source. The same dependence on the previous pulse duration and on the pulse duration for the prediction of pulse onset and offset, respectively, were observed with the subset of ORNs stimulated with pheromone loads (10^-5^ and 10^-4^ ng, see Materials and Methods) that can be directly compared with the one used to stimulate PNs (Fig 8G-H). For PNs the prediction of puff onsets was as high for isolated puffs (time interval > 1.6 s) as for puffs immediately following another one (time interval < 0.1 s), and was even larger for puffs with intermediate time intervals (between 0.4 and 0.8 s; Fig 8G). Moreover, puff onsets were always better predicted for PNs than for ORNs and the optimal prediction of puff offset in PNs was extended to puffs lasting up to 0.8 s (Fig 8H).

We repeated this analysis with the subsets of ORN data stimulated with the same pheromone load and with subsets of ORN and PN data stimulated with sequences simulating the same distance (Fig 9). In ORNs the prediction of puff onsets decreased when lowering the pheromone load (ANOVA, p < 10^-10^) independently of the duration of the preceding blanks (Fig 9A, middle panel). Puff onset prediction did not depend on the distance (Fig 9B, middle panel; ANOVA, p = 0.22). The coding of the offset of puffs in ORNs also depended on the pheromone load (Fig 9A, right panel; ANOVA, p<0.01) and was more accurate at long distances than at short ones (Fig 9B, right panel; ANOVA, p < 10^-10^). Similarly, if analyzing only ORNs stimulated with the pheromone loads (10^•5^ and 10^-4^ ng) that can be directly compared with the one used to stimulate PNs, ρ depended on distance for puff offsets (Fig 9C, right panel; ANOVA, p<0.01) but not for puff onsets (Fig 9C, middle panel; ANOVA, p = 0.56). For PNs, correlation coefficients were independent of the distance both during the onset of puffs (ANOVA, p = 0.66) and the offset of puffs (ANOVA, p = 0.75; Fig 9D). It then appears that PNs encoded efficiently the beginning and end of puffs over a broad range of temporal scales, irrespective of the temporal dynamic of the stimulation sequence.

**Figure 9.**
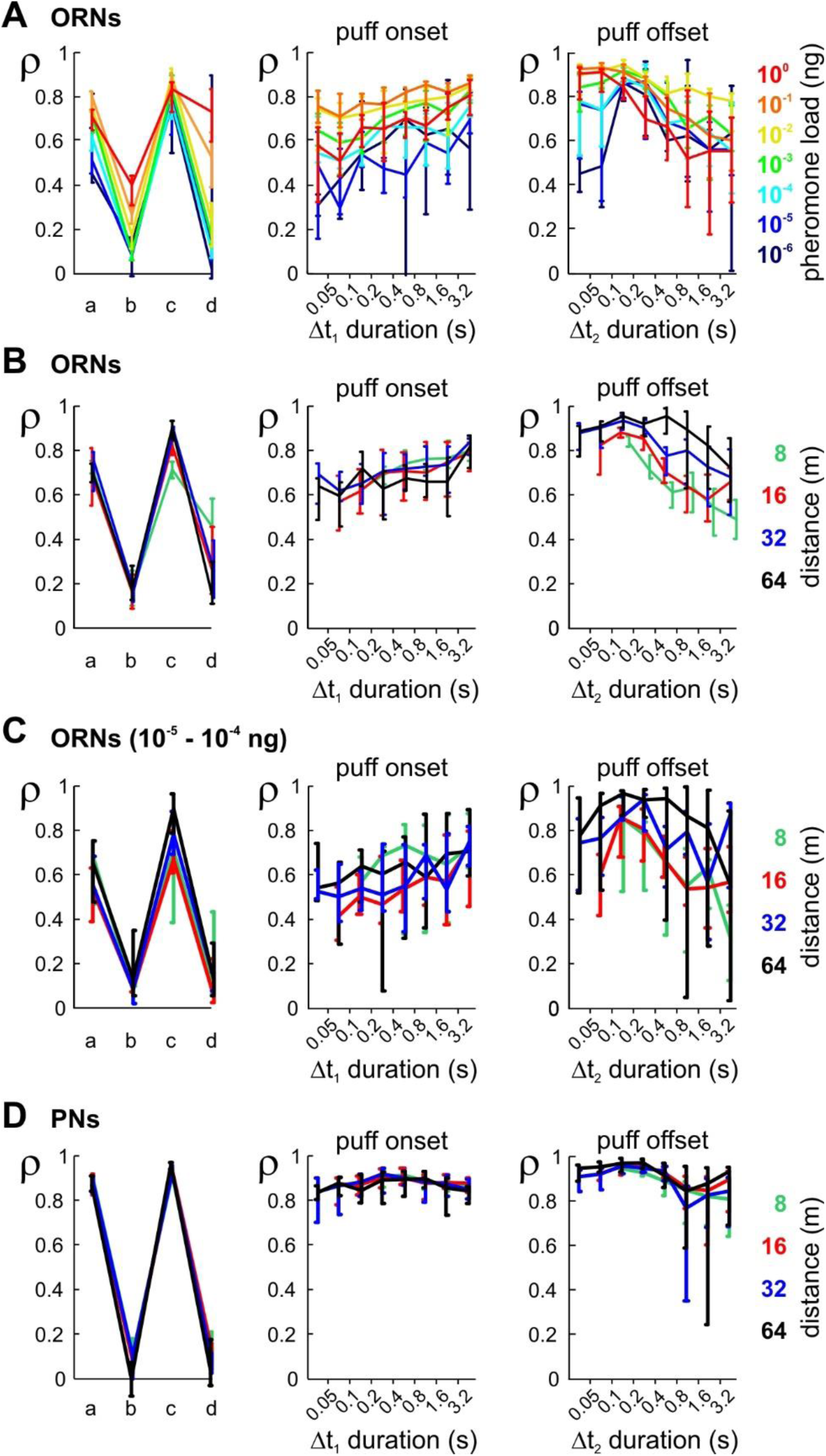
Neuronal coding of time sub-regions of turbulent stimuli analyzed as function of pheromone load and simulated distance. (A) Analysis of olfactory coding in ORNs as a function of pheromone load. Left: coding during the 4 sub-regions described in Fig 8A analyzed for each pheromone load using the same analysis as in Fig 8E. Plot shows median values of correlation coefficients and bootstrap confidence interval. Colors code for pheromone load. Middle and right: same analysis as in Fig 8G and H, but ORN data are subdivided in different pheromone loads. Middle: coding of onset of puffs depends on time delay from the end of the preceding puff and on pheromone load. Right: coding of offset of puffs depends on time delay from the end of the preceding puff and on pheromone load. (B) Sub-region analysis of olfactory coding in ORNs as a function of distance. Same conventions as in (A), but ORN data are subdivided in different simulated distances. (C) Same analysis as in (B) but from the subpopulation of ORNs stimulated with pheromone loads of 10^-5^ and 10^-4^ ng. (D) Sub-region analysis of olfactory coding in PNs as a function of distance. Same conventions as in (A), but PN data are subdivided in different distances.

## Discussion

### ORN and PN coding of broadband temporal sequences

Single-unit neuronal firing recorded from ORNs and PNs in response to sequences with white-noise and turbulent temporal statistics were analyzed with L-N models. L-N models were fitted with white-noise stimuli and used for predicting responses to turbulent stimuli. L-N models have proven to successfully predict the response of ORNs [1, 49, 50] and PNs [46] to time-varying olfactory stimuli. In addition, L-N models revealed correlations between timevarying chemosensory signals and orientation behavior both in *C. elegans* [51] and *Drosophila* larva [52-54]. Surprisingly, our study demonstrates that L-N modeling is also suited to predict neuronal responses to odorant stimuli with long-tail distributions expected from turbulent transport. As for many sensory systems, the L-N model is unable to predict adequately neuronal responses during long odor puffs but the coding of specific features is preserved, namely the timing of beginning and end of puffs.

ORN responses are generated locally, by detection of odorants and transduction into trains of action potentials. Olfactory detection depends both on pre-neuronal filtering performed by the sensillum cuticle and lymph and on the ORN intrinsic properties [55]. Olfactory information in PNs is inherited from ORN inputs. However, the processing is also refined by intrinsic properties of projection neurons (PNs) and a neuronal network including local neurons (LNs), PNs and modulatory connections from other brain areas [56-58]. We noticed significant differences in the coding properties of turbulent stimuli between ORNs and PNs, especially at large distances from the odor source. The variability of linear filters was higher in PNs than in ORNs. PNs also coded more adequately for the transition between long and short puffs than ORNs. This phenomenon was due to the inhibition phase at the end of PN responses that contrasted with the adapted response and preserved its detectability. In addition, the firing rate was way below the spontaneous activity during the PN inhibition phase, while in ORNs the spontaneous activity was already close to 0. In our model, the inhibition phase was included in the shape of the linear filters, contributing to the difference between ORN and PN filters, in particular in the PC2 component. The inhibition phase results either from local inhibitory circuits [58-60] and/or from intrinsic PN properties [61]. It is noteworthy that inhibitory mechanisms have been hypothesized to enlarge the temporal bandwidth transmitted through ORN to PN synapses [62]. Previous studies reported that ensemble of PNs can very reliably track temporal properties of the stimulus [63]. Here we show that single PNs can encode puff onset and puff offset for a large diversity of temporal structures of stimuli, and thus virtually at any distance from the source, a key information component moths can use for localizing odor sources from large distances.

The statistic of turbulent olfactory signals changes with distance from the source. Adaptation of neural coding to input signal statistics is found in numerous sensory systems including the visual [64, 65], the auditory [66-69], the mechanosensory [70, 71] and the somatosensory systems [72, 73]. Adaptation is not solely desensitization, but it changes how sensory systems process information. Here, we showed that rapid adaptation does not preclude the detection of plume transitions in PNs. Other stimulus properties could still be encoded in the PN adapted response, like odor identity through synchronous activity [74]. We also showed that a slow adaptation occurs both in ORNs and PNs, at the scale of tens of seconds. Growing evidence indicates that adaptation operates at multiple scales and needs to be implemented in L-N models in order to fully describe the neuronal dynamic [75-80]. In natural conditions ORNs and PNs are likely never fully adapted but rather oscillate dynamically between multiple adaptation levels.

### Toward more realistic stimuli

Analyzing the neuronal coding of turbulent stimuli requires a mean to monitor precisely the time course of olfactory plumes. Two strategies have been used so far: the use of biological (electroantennographic, EAG) [3, 74] or artificial odor detectors [1, 2]. However, both methods are restricted to odor detections over short distances, up to a couple of meters. Moreover, artificial detectors target a specific range of chemicals and EAG signals already result from a neuronal processing of olfactory information. Finally, it is neither possible to precisely control and set the stimulation, nor to ascertain the relative position of the recording site to the center of the plume cone. We chose to overcome these limitations by using an alternative strategy: we delivered olfactory sequences with plume time statistics corresponding to turbulent flows and tested numerous concentrations and long distance transportation. Significant work is still necessary to complete our understanding of the key features extracted by olfactory coding. Fluctuations of concentration with distance and time will also have to be taken into account. The features used by moths to provide information about source position remain unknown. Thus, a question remains on how many features present in real turbulent flows should be simulated in virtual environments. For example, in the current simulations, duration of puffs and blanks were generated to mimic turbulence structures. Yet, finer structures exist in plume dynamics [8]. In long puffs, groups of high concentration peaks are grouped together (clumps). There are relations between the statistics of peak numbers and the distance to the source, thus moths might be able to exploit these peaks to estimate the distance to the source.

The dynamic of odorant reaching the insect antenna depends not only on the turbulence but also on active olfactory sampling behaviors including wing beating, antennal flicking and body displacement [74, 81, 82]. In realistic olfactory searches strong correlations in detection dynamics (time and concentration) are due to the strategy implemented by moths. Thus, a complete study would require effective modelling of the moth search strategies. However, long distance recording of searching moths coupled to the olfactory input is not accessible. Thus, it is unknown whether moths or other insects are really able to perform olfactory searches at very large distances solely based on the specific input of the odor of interest. Nevertheless, our study unveils the basic mechanisms of olfactory processing of the broadband temporal profiles of olfactory plumes in the turbulent realm.

## Materials and Methods

### Insects

*A. ipsilon* moths were reared on an artificial diet until pupation. Adults were maintained under a reversed 16h:8h light:dark photoperiod at 23°C and fed with sugar water. Experiments were performed on virgin 4- or 5-day-old (sexually mature) males.

### Electrophysiology

For all recordings, insects were immobilized with the head protruding. For ORN recordings, the antenna was fixed with adhesive tape on a small support and a tungsten electrode was inserted at the base of a pheromone-sensitive sensillum. The electrical signal was amplified (×1000) and band-pass filtered (10 Hz – 5 kHz) with an ELC-03X (npi electronic, Tamm, Germany), and digitized at 10 kHz (NI-9215, National Inst., Nanterre, France) under Labview (National Inst.).

For PN recordings, the head was immobilized with dental wax after removing the scales. The cephalic capsule was open and the brain was exposed by removing the mouth parts and muscles. The antennal lobe was desheathed to allow the penetration of the recording electrode, a glass micropipette whose tip was manually broken to a diameter of 2 µm and filled with (in mM): NaCl 150, KCl 4, CaCl_2_ 6, MgCl_2_ 2, Hepes 10, Glucose 5 (pH 7.2). The same solution was used to perfuse the preparation. The pipette was slowly inserted in the cumulus of the macroglomerulus of one antennal lobe, where ORNs sensitive to the major pheromone component project [45], until the appearance of a well-isolated single-unit firing activity. Extracellular recordings from *A. ipsilon* proved to sample only neurons with a large neurite (PNs but not LNs) in the antennal lobe [45]. Furthermore, the inhibition phase recorded at the end of each individual response is a hallmark of PN responses both in this species [83] and in other insect species such as *Manduca sexta* [84] and *D. melanogaster* [85]. The electrical signal was amplified (×1000) and band-pass filtered (10 Hz – 5 kHz) with a Cyberamp 320 (Molecular Devices, Union City, CA, USA) and digitized at 10 KHz (NI-9215) under Labview.

### Stimulation

We used the major component, (*Z*)-7-dodecenyl acetate (Z7-12:Ac) of the *A. ipsilon* sex pheromone blend as stimulus. Z7-12:Ac was diluted in hexane and applied to a filter paper at doses ranging from 10^-6^ to 10^0^ ng (ORN recordings) and at 10^-1^ ng (PN recordings). The entire antenna was exposed to a constant charcoal-filtered and humidified air flow (70 L.h^-1^). For ORN recordings, air puffs (10 L.h^-1^) were delivered through a calibrated capillary (Ref. 11762313, Fisher Scientific, France) positioned at 1 mm from the antenna and containing the odor-loaded filter paper (10 × 2 mm). For PN recordings, stimuli were applied by inserting 15 cm upstream of a 7-mm glass tube a Pasteur pipette containing the odor-loaded filter paper (15 × 10 mm). Responses to a control (hexane) and pheromone puffs of increasing concentrations (10^-3^ to 10^1^ ng) were first tested. Only PNs exhibiting dose-dependent pheromone responses were included in the analysis.

Control of electrovalves (LHDA-1233215-H, Lee Company, France) was done by custom-made Labview programs. All sequences were executed asynchronously from text files where the states of electrovalves were indicated by a Boolean variable. Sequences were generated using Matlab scripts (The Mathworks, Natick, MA). The time resolution of the sequences was 1 ms. The characteristic response time of the valves, *i.e.* the time to go from open to closed (and closed to open) is < 5 ms.

### Stimulus Sequences

White-noise stimuli were approximated by randomly controlling the state (Open-Close) of the electrovalve used to deliver pheromone stimuli. The valve state was updated every 10, 50 or 100 ms (referred to the correlation time). Thus, the spectra were colored for frequencies greater than 16, 3.4 and 1.6 Hz, respectively. There is no significant effect of correlation times on filter extraction.

Time dynamics of turbulent plumes were derived from Celani et al. [8]. The virtual source and positions of the moths were aligned with the wind direction. We tested 4 virtual distances, 8, 16, 32, 64 m. The geometric progression of distances was chosen to emphasize the effect of turbulence on puff/blank statistics. The distribution of puff/blank time was generated from:

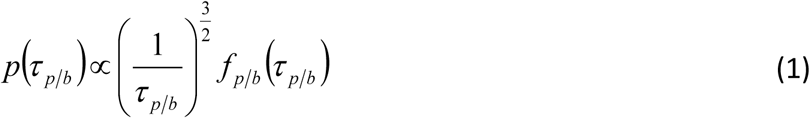

where the index *p* is for puff, *b* for blanks, *f*_*p/b*_ is a cutoff function with exponential decrease of rate 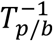 for τ_*p/b*_≥*T*_*p/b*_. Cutoffs *T*_*p/b*_ are physical properties of the turbulent flow. Using similar notation as in Celani et al. [8], we set *U* = 1 m.s^-1^ (average wind velocity), *δU* = 0.1 m.s^-1^ (wind fluctuations), *a* = 0.1 m (size of the pheromone source), *χ*= 0.4 (intermittency factor), 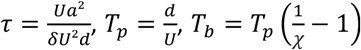. The largest distance used for ORNs, 64 m, imposed a selection on the generated sequences because some sequences exhibited either extremely long stimuli, which led to the complete stop of spiking activity, or almost no puffs. Thus, the statistics for 64 m was biased from the pure turbulence by removal of extremely rare events (puffs > 30 s). A given sequence of turbulent olfactory sequence was never used twice with the same pheromone concentration unless specifically mentioned for tests with frozen sequences.

A single ORN was recorded per insect. For each recorded ORN, only one sequence of stimuli was delivered during 15 min at a constant pheromone concentration. The different stimulation sequences were tested on different ORNs. White-noise stimulations were performed with 3 correlation times (10, 50, 100 ms) and 4 decades of pheromone concentrations. Turbulent plumes were simulated from 4 distances and 7 decades of concentrations. Control experiments were performed with 2 repetitions of 3-min long white-noise sequences interleaved with various (1, 3, 5, 15 min) inter-sequence intervals.

We tested PNs at 0.1 ng of pheromone loaded on the filter paper, a dose chosen in the medium-low range of cell population dose-response curves [45]. Stimuli were delivered to the entire antenna to ensure proper coding in PNs. For each recorded neuron several stimulation sequences were delivered to the antenna. The first sequence was a 4-min long white-noise stimulation followed by 4 min of no stimuli. The white-noise sequence had a correlation time of 50 ms. Then 4 different sequences of olfactory plume statistics simulating 4 distances were presented in random order. Sequence durations were 5 min and inter-sequence intervals were 3 min. Control experiments were performed in the following way: (1) delivering 4 consecutive frozen white-noise sequences with various inter-sequence intervals, (2) delivering 3 consecutive frozen sequences with a temporal dynamic simulating a distance of 16 m and (3) testing 3 white-noise sequences with different correlation times, 10, 50 and 100 ms.

### Data Analysis

Spike sorting was performed with Spike2 (CED, Oxford). Shapes of action potentials were analyzed using principal component analysis and spike clusters were identified using the first 3 eigenvector subspaces. In long duration experiments involving turbulent stimuli, slow drifts of clusters and secondary clusters resulting from collision with spikes from another cell were taken into account. Finally, visual inspection was used to ensure that rare events with high frequency did not prevent efficient spike detections. Firing rates were calculated by Gaussian convolution of 50 ms width then resampled at 100 Hz.

### L-N Model

To model neural responses we used the linear–nonlinear (L-N) model. The L-N model was determined separately for each neuron, and for each dose as one single dose of pheromone was applied on each neuron. The firing rate *r*(*t*) of neurons is decomposed into a linear kernel *K*(*t*) and a nonlinear function *H*(*r*). Thus we have

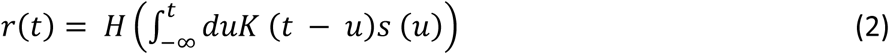

with *s*(*t*) the olfactory input signal. The kernel is extracted by minimizing the L_2_ norm between experimental measures and the L-N prediction with a Lasso regularization [86]

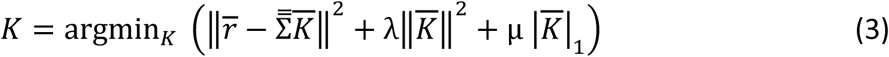

with 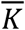 the Kernel Vector, 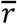 the experimental vector response, 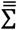 the time shifted stimulus matrix and (λ, µ) the regularization coefficients. Lasso regularization was used to prevent anomalies in *K*(*t*) due both to overfitting and to correlations in the input signal *s*(*t*). The L_2_ norm prevents anomalous rise of the Kernel amplitude and L_1_ norm favors sparse response. The nonlinear function *H*(*r*) was extracted by fitting the experimental rate, *r*(*t*), as a function of the linear rate,

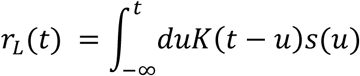

using a Hill function. The training samples were made by 20% of the points and the test sample by the remaining 80%. The procedure was applied 100 times per sequence to set the (λ, µ) values.

### Modified L-N Model

In order to sample coding for turbulent environments, recordings were performed over long time periods (up to 15 min). The firing rate was found to decrease during a stimulus sequence in a predictable and reversible fashion. Thus the L-N model with a short-term kernel was completed adding a long-term kernel leading to the effective *K*_*all*_(*t*) = *K*_*short*_(*t*) + *βK*_*long*_(*t*). The kernel was extracted from equation 2 where the time shifted stimulus matrix was completed with 7 additional columns corresponding to the long-time kernel. The columns contained the integrated firing rate calculated in time windows with respective edges of 2.5, 5, 10, 20, 40, 80, 160 and 320 s.

The accuracy of the filters was quantified using classical *R*^2^ determination coefficients between prediction and measured data. *R*^2^ is the proportion of variance in the measured firing rate (independent variable) that is predicted by the model (dependent variable). For data vs data comparison, as well as predicted vs measured data in small time regions, both variables were relatively independent with different mean signals, and thus correlation coefficients, *ρ*, were used since more informative than *R*^2^.

### Comparison of odor delivery devices

As we used different odor delivery devices when recording ORNs (peri-sensillum device) and PNs (whole-antenna device), we established dose-response curves from ORNs with both devices to compare them (200-ms stimuli; n = 7). We measured the maximum firing frequency during the response and the number of action potentials evoked until the firing frequency returned to baseline. The comparison of pheromone loads and responses indicates that the concentration delivered on a sensillum by the peri-sensillum device is 10^3^ to 10^4^ higher than with the whole-antenna stimulator (Fig S1A). We inferred that 0.1 ng stimuli with the whole-antenna stimulator correspond approximately to 10^-5^ to 10^-4^ ng stimuli with the peri-sensillum device. ORN activities (n = 32) were then recorded in response to a frozen white-noise sequence (5 min, time step = 50 ms) delivered consecutively by both devices (10 ng with the whole antenna device and 5 pg with the peri-sensillum device). The order of use of the two devices was randomized. The average short-term filters calculated from the responses to the white-noise sequences delivered by the two devices overlap (Fig S1B), indicating that they deliver stimuli with the same dynamics.

## Acknowledgments

This work was funded by the state program Investissements d’avenir managed by ANR (grant ANR-10-BINF-05 “Pherotaxis”). V.J. was also funded by the Conseil Régional de la Réunion and by the European regional development fund (ERDF). The authors are thankful to Massimo Vergassola and Antonio Celani for helpful discussions and to Sylvia Anton, Taro Toyoizumi and Jürgen Reingruber for their critical reading of the manuscript.

---

**Figure S1.**
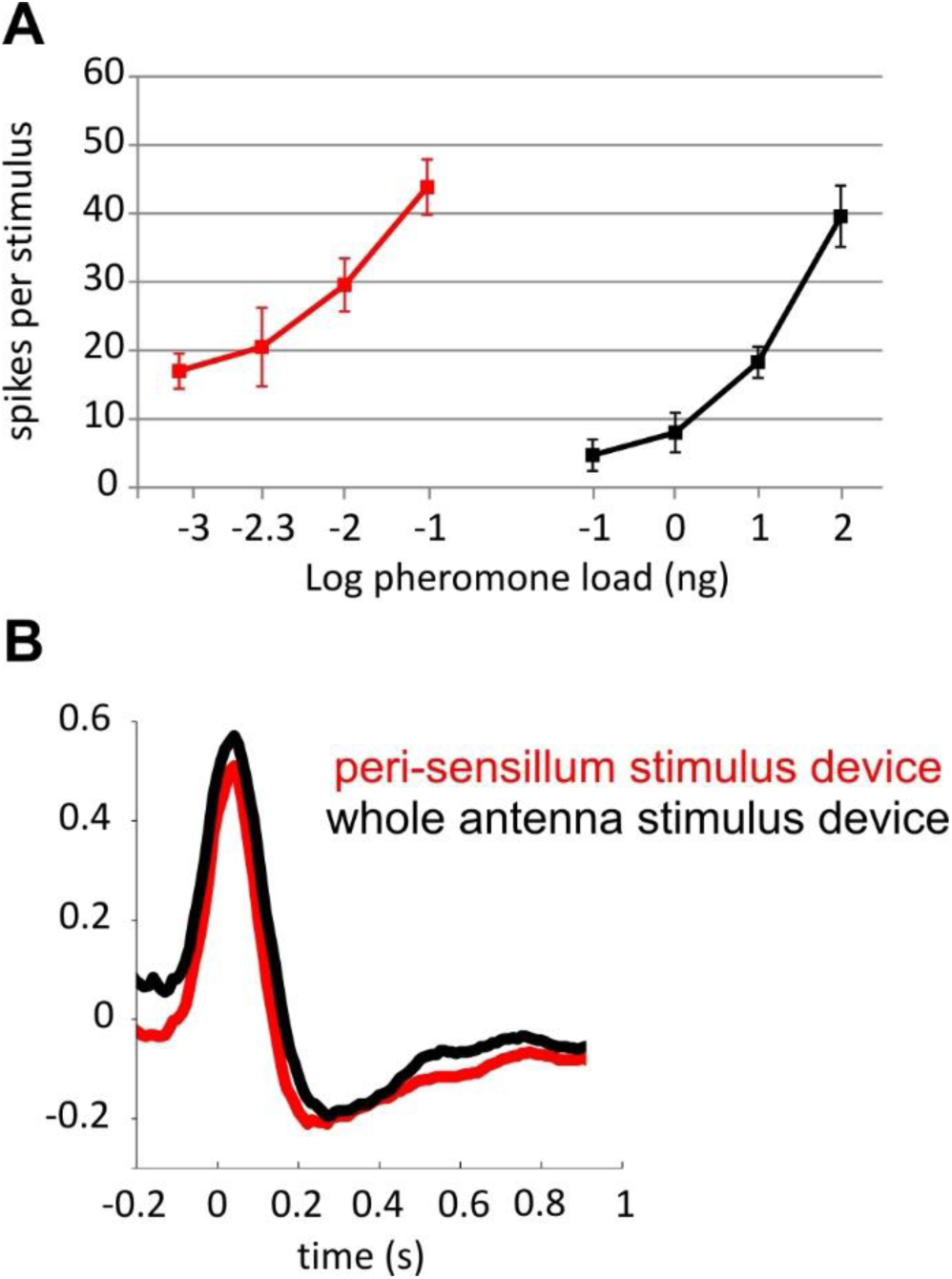
Comparison of whole antenna and peri-sensillum odor delivery devices. (A) Number of action potentials evoked until the firing frequency returned to baseline in response to 200-ms stimuli with different pheromone loads (n = 7). The 10 ng pheromone load used with the whole antenna stimulator induced approximatively the same ORN responses as 1 to 10 pg used with the peri-sensillum stimulator. (B) Average short-term filters calculated from ORN responses to a frozen noise sequence (time step = 50 ms) delivered consecutively by both stimulus devices while an ORN activity was recorded (n = 32). The pheromone load was 10 ng with the whole antenna stimulator and 5 pg with the peri-sensillum stimulator. The order of use of the two stimulators was randomized. The 2 linear filters were aligned on the peak of the positive lobe to compensate for the longer delay in odor delivery with the whole antenna device.

## REFERENCES

1. Nagel KI, Wilson RI. Biophysical mechanisms underlying olfactory receptor neuron dynamics. Nat Neurosci. 2011;14(2):208–16.

2. Riffell JA, Shlizerman E, Sanders E, Abrell L, Medina B, Hinterwirth AJ, et al. Flower discrimination by pollinators in a dynamic chemical environment. Science. 2014;344(6191):1515–8.

3. Vickers NJ, Christensen TA, Baker TC, Hildebrand JG. Odour-plume dynamics influence the brain’s olfactory code. Nature. 2001;410(6827):466–70.

4. Gorur-Shandilya S, Demir M, Long J, Clark DA, Emonet T. Olfactory receptor neurons use gain control and complementary kinetics to encode intermittent odorant stimuli. Elife. 2017;6:e27670.

5. Falkovich G, Gawedzki. K, Vergassola M. Particles and fields in fluid turbulence. Rev Mod Phys. 2001;73(4):913–75.

6. Murlis J, Elkinton JS, Cardé RT. Odor plumes and how insects use them. Annu Rev Entomol. 1992;37:505–32.

7. Weissburg MJ. The fluid dynamical context of chemosensory behavior. Biol Bull. 2000;198(2):188–202.

8. Celani A, Villermaux E, Vergassola M. Odor landscapes in turbulent environments. Phys Rev X. 2014;4:041015.

9. Cardé RT, Willis MA. Navigational strategies used by insects to find distant, wind-borne sources of odor. J Chem Ecol. 2008;34(7):854–66.

10. Vickers NJ. Winging it: moth flight behavior and responses of olfactory neurons are shaped by pheromone plume dynamics. Chem Senses. 2006;31(2):155–66.

11. Cardé RT, Charlton RE. Olfactory sexual communication in Lepidoptera: strategy, sensitivity and selectivity. In: Lewis T, editor. Insect communication: Academic press, London; 1984. pp. 241–65.

12. Elkinton JS, Schal C, Ono T, Cardé RT. Pheromone puff trajectory and upwind flight of male gypsy moths in a forest. Physiol Entomol. 1987;12:399–406.

13. Shorey HH. Animal Communication by Pheromones: Academic Press Inc.; 1976. 175 p.

14. Rebello MR, McTavish TS, Willhite DC, Short SM, Shepherd GM, Verhagen JV. Perception of odors linked to precise timing in the olfactory system. PLoS Biol. 2014;12(12):e1002021.

15. Shusterman R, Smear MC, Koulakov AA, Rinberg D. Precise olfactory responses tile the sniff cycle. Nat Neurosci. 2011;14(8):1039–44.

16. Smear M, Shusterman R, O’Connor R, Bozza T, Rinberg D. Perception of sniff phase in mouse olfaction. Nature. 2011;479(7373):397–400.

17. Smear M, Resulaj A, Zhang J, Bozza T, Rinberg D. Multiple perceptible signals from a single olfactory glomerulus. Nat Neurosci. 2013;16(11):1687–91.

18. Carey RM, Verhagen JV, Wesson DW, Pirez N, Wachowiak M. Temporal structure of receptor neuron input to the olfactory bulb imaged in behaving rats. J Neurophysiol. 2009;101(2):1073–88.

19. Gupta P, Albeanu DF, Bhalla US. Olfactory bulb coding of odors, mixtures and sniffs is a linear sum of odor time profiles. Nat Neurosci. 2015;18(2):272–81.

20. Scott JW. Sniffing and spatiotemporal coding in olfaction. Chem Senses. 2006;31(2):119–30.

21. Verhagen JV, Wesson DW, Netoff TI, White JA, Wachowiak M. Sniffing controls an adaptive filter of sensory input to the olfactory bulb. Nat Neurosci. 2007;10(5):631–9.

22. Mozell MM, Jagodowicz M. Chromatographic separation of odorants by the nose: retention times measured across *in vivo* olfactory mucosa. Science. 1973;181(4106):1247–9.

23. Scott JW, Brierley T, Schmidt FH. Chemical determinants of the rat electro-olfactogram. J Neurosci. 2000;20(12):4721–31.

24. Ezeh PI, Davis LM, Scott JW. Regional distribution of rat electroolfactogram. J Neurophysiol. 1995;73(6):2207–20.

25. Kent PF, Mozell MM, Murphy SJ, Hornung DE. The interaction of imposed and inherent olfactory mucosal activity patterns and their composite representation in a mammalian species using voltage-sensitive dyes. J Neurosci. 1996;16(1):345–53.

26. Willis MA, Ford EA, Avondet JL. Odor tracking flight of male *Manduca sexta* moths along plumes of different cross-sectional area. J Comp Physiol A. 2013;199(11):1015–36.

27. Kennedy JS. Zigzagging and casting as a preprogrammed response to wind-borne odour: A review. Physiol Entomol. 1983;27:58–66.

28. Kramer E. A tentative intercausal nexus and its computer model on insect orientation in windborne pheromone plumes. In: Cardé RT, Minks AK, editors. Insect pheromone research: new directions. Kluwer Academic Publishers ed; 1997. pp. 232–48.

29. Vickers NJ, Baker TC. Reiterative responses to single strands of odor promote sustained upwind flight and odor source location by moths. Proc Natl Acad Sci USA. 1994;91(13):5756–60.

30. Justus KA, Cardé RT. Flight behaviour of males of two moths, *Cadra cautella* and *Pectinophora gossypiella*, in homogeneous clouds of pheromone. Physiol Entomol. 2002;27:67–75.

31. Kennedy JS, Ludlow AR, Sanders CJ. Guidance system used in moth sex attraction. Nature. 1980;288:475–7.

32. Willis MA, Baker TC. Effect of intermittent and continuous pheromone stimulation on the flight behaviour of the oriental fruit moth, *Grapholita molesta*. Physiol Entomol. 1984;9:341–58.

33. Kramer E. Attractivity of pheromone is surpassed by time-patterned application of two nonpheromone compounds. J Insect Behav. 1992;5(1):83–97.

34. Baker TC, Willis MA, Haynes KF, Phelan PL. A pulsed cloud of pheromone elicits upwind flight in male moths. Physiol Entomol. 1985;10:257–65.

35. Mafra-Neto A, Cardé RT. Fine-scale structure of pheromone plumes modulates upwind orientation of flying moths. Nature. 1994;369(6476):142–4.

36. Vickers NJ, Baker TC. Male *Heliothis virescens* maintain upwind flight in response to experimentally pulsed filaments of their sex pheromone (Lepidoptera: noctuidae). J Insect Behav. 1992;5:669–87.

37. Willis MA, David CT, Murlis J, Cardé RT. Effects of pheromone plume structure and visual stimuli on the pheromone-modulated upwind flight of male gypsy moths (*Lymantria dispar*) in a forest (Lepidoptera: Lymantriidae). J Insect Behav. 1994;7:385–409.

38. Mafra-Neto A, Cardé RT. Influence of plume structure and pheromone concentration on upwind flight of *Cadra cautella* males. Physiol Entomol. 1995;20:117–33.

39. Barlow HB. Possible principles underlying the transformations of sensory messages. In: Rosenblith WA, editor. Sensory Communication: Cambridge, MA: MIT Press; 1961. pp. 217–34.

40. Fairhall AL, Lewen GD, Bialek W, de Ruyter Van Steveninck RR. Efficiency and ambiguity in an adaptive neural code. Nature. 2001;412(6849):787–92.

41. Kostal L, Lansky P, Rospars JP. Efficient olfactory coding in the pheromone receptor neuron of a moth. PLoS Comput Biol. 2008;4(4):e1000053.

42. Simoncelli EP, Olshausen BA. Natural image statistics and neural representation. Annu Rev Neurosci. 2001;24:1193–216.

43. Dong D, Atick J. Statistics of natural time-varying images. Network: Comput Neural Syst 1995;6:345–58.

44. Justus KA, Murlis J, Jones C, Cardé RT. Measurement of odor-plume structure in a wind tunnel using a photoionization detector and a tracer gas. Environ Fluid Mech. 2002;2:115–42.

45. Rospars JP, Gremiaux A, Jarriault D, Chaffiol A, Monsempes C, Deisig N, et al. Heterogeneity and convergence of olfactory first-order neurons account for the high speed and sensitivity of second-order neurons. PLoS Comput Biol. 2014;10(12):e1003975.

46. Geffen MN, Broome BM, Laurent G, Meister M. Neural encoding of rapidly fluctuating odors. Neuron. 2009;61(4):570–86.

47. Bussgang JJ. Cross-correlation functions of amplitude-distorted Gaussian signals. MIT Research Laboratory Technical Report. 1952; no 216.

48. Murlis J, Jones CD. Fine-scale structure of odour plumes in relation to insect orientation to distant pheromone and other attractant source. Physiol Entomol. 1981;6:71–86.

49. Kim AJ, Lazar AA, Slutskiy YB. System identification of *Drosophila* olfactory sensory neurons. J Comput Neurosci. 2011;30(1):143–61.

50. Martelli C, Carlson JR, Emonet T. Intensity invariant dynamics and odor-specific latencies in olfactory receptor neuron response. J Neurosci. 2013;33(15):6285–97.

51. Kato S, Xu Y, Cho CE, Abbott LF, Bargmann CI. Temporal responses of *C. elegans* chemosensory neurons are preserved in behavioral dynamics. Neuron. 2014;81(3):616–28.

52. Gepner R, Mihovilovic Skanata M, Bernat NM, Kaplow M, Gershow M. Computations underlying *Drosophila* photo-taxis, odor-taxis, and multi-sensory integration. Elife. 2015;4:06229.

53. Hernandez-Nunez L, Belina J, Klein M, Si G, Claus L, Carlson JR, et al. Reverse-correlation analysis of navigation dynamics in *Drosophila* larva using optogenetics. Elife. 2015;4:e06225.

54. Schulze A, Gomez-Marin A, Rajendran VG, Lott G, Musy M, Ahammad P, et al. Dynamical feature extraction at the sensory periphery guides chemotaxis. Elife. 2015;4:e06694.

55. Suh E, Bohbot J, Zwiebel LJ. Peripheral olfactory signaling in insects. Curr Opin Insect Sci. 2014;6:86–92.

56. Hu A, Zhang W, Wang Z. Functional feedback from mushroom bodies to antennal lobes in the *Drosophila* olfactory pathway. Proc Natl Acad Sci USA. 2010;107(22):10262–7.

57. Masse NY, Turner GC, Jefferis GS. Olfactory information processing in *Drosophila*. Curr Biol. 2009;19(16):R700–R13.

58. Wilson RI. Early olfactory processing in *Drosophila*: mechanisms and principles. Annu Rev Neurosci. 2013;36:217–41.

59. Christensen TA, Waldrop BR, Hildebrand JG. Multitasking in the olfactory system: context-dependent responses to odors reveal dual GABA-regulated coding mechanisms in single olfactory projection neurons. J Neurosci. 1998;18(15):5999–6008.

60. Wilson RI, Laurent G. Role of GABAergic inhibition in shaping odor-evoked spatiotemporal patterns in the *Drosophila* antennal lobe. J Neurosci. 2005;25(40):9069–79.

61. Martinez D, Chaffiol A, Voges N, Gu Y, Anton S, Rospars JP, et al. Multiphasic on/off pheromone signalling in moths as neural correlates of a search strategy. PLoS One. 2013;8(4):e61220.

62. Nagel KI, Hong EJ, Wilson RI. Synaptic and circuit mechanisms promoting broadband transmission of olfactory stimulus dynamics. Nat Neurosci. 2015;18(1):56–65.

63. Brown SL, Joseph J, Stopfer M. Encoding a temporally structured stimulus with a temporally structured neural representation. Nat Neurosci. 2005;8(11):1568–76.

64. Brenner N, Bialek W, de Ruyter van Steveninck R. Adaptive rescaling maximizes information transmission. Neuron. 2000;26(3):695–702.

65. Lesica NA, Jin J, Weng C, Yeh CI, Butts DA, Stanley GB, et al. Adaptation to stimulus contrast and correlations during natural visual stimulation. Neuron. 2007;55(3):479–91.

66. Lewicki MS. Efficient coding of natural sounds. Nat Neurosci. 2002;5(4):356–63.

67. Dahmen JC, Keating P, Nodal FR, Schulz AL, King AJ. Adaptation to stimulus statistics in the perception and neural representation of auditory space. Neuron. 2010;66(6):937–48.

68. Dean I, Harper NS, McAlpine D. Neural population coding of sound level adapts to stimulus statistics. Nat Neurosci. 2005;8(12):1684–9.

69. Watkins PV, Barbour DL. Specialized neuronal adaptation for preserving input sensitivity. Nat Neurosci. 2008;11(11):1259–61.

70. Clague H, Theunissen F, Miller JP. Effects of adaptation on neural coding by primary sensory interneurons in the cricket cercal system. J Neurophysiol. 1997;77(1):207–20.

71. Zheng HJ, Wang Q, Stanley GB. Adaptive shaping of cortical response selectivity in the vibrissa pathway. J Neurophysiol. 2015;113(10):3850–65.

72. Garcia-Lazaro JA, Ho SS, Nair A, Schnupp JW. Shifting and scaling adaptation to dynamic stimuli in somatosensory cortex. Eur J Neurosci. 2007;26(8):2359–68.

73. Maravall M, Petersen RS, Fairhall AL, Arabzadeh E, Diamond ME. Shifts in coding properties and maintenance of information transmission during adaptation in barrel cortex. PLoS Biol. 2007;5(2):e19.

74. Huston SJ, Stopfer M, Cassenaer S, Aldworth ZN, Laurent G. Neural encoding of odors during active sampling and in turbulent plumes. Neuron. 2015;88(2):403–18.

75. Drew PJ, Abbott LF. Models and properties of power-law adaptation in neural systems. J Neurophysiol. 2006;96(2):826–33.

76. Lundstrom BN, Fairhall AL, Maravall M. Multiple timescale encoding of slowly varying whisker stimulus envelope in cortical and thalamic neurons *in vivo*. J Neurosci. 2010;30(14):5071–7.

77. Lundstrom BN, Higgs MH, Spain WJ, Fairhall AL. Fractional differentiation by neocortical pyramidal neurons. Nat Neurosci. 2008;11(11):1335–42.

78. Pozzorini C, Naud R, Mensi S, Gerstner W. Temporal whitening by power-law adaptation in neocortical neurons. Nat Neurosci. 2013;16(7):942–8.

79. Sharpee TO, Sugihara H, Kurgansky AV, Rebrik SP, Stryker MP, Miller KD. Adaptive filtering enhances information transmission in visual cortex. Nature. 2006;439(7079):936–42.

80. Willmore BD, Schoppe O, King AJ, Schnupp JW, Harper NS. Incorporating midbrain adaptation to mean sound level improves models of auditory cortical processing. J Neurosci. 2016;36(2):280–9.

81. Houot B, Burkland R, Tripathy S, Daly KC. Antennal lobe representations are optimized when olfactory stimuli are periodically structured to simulate natural wing beat effects. Front Cell Neurosci. 2014;8:159.

82. Tabuchi M, Sakurai T, Mitsuno H, Namiki S, Minegishi R, Shiotsuki T, et al. Pheromone responsiveness threshold depends on temporal integration by antennal lobe projection neurons. Proc Natl Acad Sci USA. 2013;110(38):15455–60.

83. Jarriault D, Gadenne C, Rospars J-P, Anton S. Quantitative analysis of sex-pheromone coding in the antennal lobe of the moth *Agrotis ipsilon*: a tool to study network plasticity. J Exp Biol. 2009;212(Pt 8):1191–201.

84. Matsumoto SG, Hildebrand JG. Olfactory mechanisms in the moth *Manduca sexta*: response characteristics and morphology of central neurons in the antennal lobes. Proc R Soc Lond B. 1981;213:249–77

85. Wilson RI, Turner GC, Laurent G. Transformation of olfactory representations in the *Drosophila* antennal lobe. Science. 2004;303(5656):366–70.

86. Zou H, Zhu J, Hastie T. New Multicategory Boosting Algorithms Based on Multicategory Fisher-Consistent Losses. Ann Appl Stat. 2008;2(4):1290–306.

